# Single-cell transcriptomics reveal hyperacute cytokine and immune checkpoint axis in patients with poor neurological outcomes after cardiac arrest

**DOI:** 10.1101/2022.04.04.487033

**Authors:** Tomoyoshi Tamura, Changde Cheng, Wenan Chen, Louis T. Merriam, Mayra Pinilla-Vera, Jack Varon, Peter C. Hou, Patrick R. Lawler, William M. Oldham, Raghu R. Seethala, Yohannes Tesfaigzi, Alexandra J. Weissman, Rebecca M. Baron, Fumito Ichinose, Katherine M. Berg, Erin A. Bohula, David A. Morrow, Xiang Chen, Edy Y. Kim, Immunology of Cardiac Arrest Network (I-CAN)

**Author notes:** Correspondence (X.C.), (E.Y.K.) Correspondence: Xiang Chen, Ph.D. 262 Danny Thomas Place MS 1135 Memphis, TN 38105, USA Edy Y. Kim, M.D., Ph.D. Hale Building for Transformative Medicine, Room 3016-I 60 Fenwood Rd., Boston, MA 02115, USA e. These authors contributed equally to this work (Tomoyoshi Tamura, Changde Cheng).

## Abstract

Neurological injury is a major driver of mortality among patients hospitalized after cardiac arrest (CA). The early systemic inflammatory response after CA is associated with neurological injury and mortality but remains poorly defined. We determine the innate immune network induced by clinical CA at single-cell resolution. Immune cell states diverge as early as 6h post-CA between patients with good or poor neurological outcomes at hospital discharge. Nectin-2^+^ monocyte and Tim-3^+^ natural killer (NK) cell subpopulations associate with poor outcomes, and interactome analysis highlights their crosstalk via cytokines and immune checkpoints. Ex vivo studies on peripheral blood cells from CA patients demonstrate that immune checkpoints are a compensatory mechanism against inflammation after CA. IFNγ/IL-10 induce Nectin-2 on monocytes; in a negative feedback loop, Nectin-2 suppresses IFNγ production by NK cells. The initial hours after CA may represent a window for therapeutic intervention in the resolution of inflammation via immune checkpoints.

## INTRODUCTION

Over 340,000 out-of-hospital cardiac arrests (OHCA) occur in the US every year, and only 10% of these patients survive ((Benjamin et al., 2019)). With improved pre-hospital resuscitation, 20- 30% of emergency medical service-treated OHCA patients survive until hospital admission, but less than 40% of hospitalized patients survive to discharge (Virani et al., 2021). Neurological injury is the primary reason for in-hospital death and causes life-altering neurological morbidity in 15-25% of patients who survive (Berdowski et al., 2010; Neumar et al., 2008).

The global ischemia-reperfusion injury (IRI) of CA leads to a profound systemic inflammatory response characterized by high levels of circulating cytokines, endotoxemia, and dysregulated immune responses (Adrie et al., 2002). Increases in proinflammatory cytokines (e.g., interleukin [IL]-6, tumor necrosis factor [TNF]-α), reduced lymphocyte counts, and high neutrophil-to-lymphocyte ratios are associated poor neurological outcome and mortality after CA (Adrie et al., 2002; Fries et al., 2009); (Miyatake et al., 2019; Weiser et al., 2017); (Villois et al., 2017)(Kim et al., 2018; Weiser et al., 2017). Transcriptomic analysis of whole blood by microarray corroborated the increased expression of proinflammatory genes is associated with worse outcomes after CA (Tissier et al., 2019). Despite prior studies suggesting that the immune response may influence neurological injury and clinical outcomes, we lack an in-depth characterization of the global immune response and its regulation after CA. In the limited studies of the immunology of clinical CA to date, the focus has been on potentially pathogenic, proinflammatory axes. The recent advance of single-cell RNA sequencing (scRNA-seq) offers unbiased profiling of the global transcriptome at single-cell resolution. scRNA-seq has identified new transcriptional cell states and previously unappreciated heterogeneity within immune cell populations (Papalexi and Satija, 2018; Reyes et al., 2020, 2021). Although chronic cardiovascular disorders, such as atherosclerotic carotid artery disease (Fernandez et al., 2019) and heart failure (Abplanalp et al., 2021) have global transcriptomes defined at single-cell resolution, no prior study has used scRNA-seq to investigate the immediate post-CA period.

Here, we characterize the immune response to clinical CA at single-cell resolution. In peripheral blood mononuclear cells (PBMC), we found that, as early as 6h post-OHCA, the global transcriptomes of innate immune cells diverged between patients with eventual poor or good neurological outcomes. *NECTIN2*^+^ monocyte and *HAVCR2*^+^ (Tim-3^+^) natural killer (NK) cell cellular states were expanded in patients with poor neurological outcomes after cardiac arrest; a finding which was confirmed at the protein level in a validation cohort of OHCA patients. In the validation cohort, we sorted for Nectin-2^+^ monocytes and Tim-3^+^ NK cells. Bulk RNA-seq of these cell subpopulations demonstrated that these immune checkpoint receptors were specific markers for the global transcriptomic cell state identified by scRNA-seq. Interactome analysis highlighted immune checkpoint receptors as a candidate axis for monocyte-NK cellular crosstalk. Ex vivo studies on PBMC from post-CA subjects established a negative feedback loop in which the mixed pro- and anti-inflammatory cytokine milieu induces expression of Nectin-2^+^ on monocytes. Nectin-2 then limited IFNγ production by NK cells (Scheme in **Figure S1**). These results reveal that CA induced inhibitory immune checkpoint receptors act as a compensatory response to ameliorate inflammation. These findings uncover a possible therapeutic window arising within the first few hours of resuscitation.

## RESULTS

### Global transcriptomic profile at single-cell resolution of PBMC post-OHCA

A total of 96,179 peripheral blood mononuclear cells (PBMC) were transcriptionally profiled at single-cell resolution from post-OHCA patients at different time points and healthy subjects.

Post-OHCA patients were categorized as having either good neurological outcomes (“good outcomes”) or poor neurological outcomes (“poor outcomes”) by Cerebral Performance Category (CPC) at hospital discharge. CPC is the most widly utilized standard for assessing neurological functional outcomes after clinical CA (Becker et al., 2011). CPC ranges from 1 to 5 and 1 and 2, and 3-5 were determined as good and poor neurological outcomes, respectively. A total of 11 OHCA patients (4 with good outcomes and 7 with poor outcomes) and 3 healthy controls were included in the discovery cohort for scRNA-seq analysis (**Figure 1A, Table S1**). Two separate validation cohorts of OHCA patients (N = 28 and 47, respectively) were enrolled. We created a browser-based visualization tool of the single-cell gene expression data to assist non-bioinformatics researchers https://viz.stjude.cloud/chen-lab/visualization/single-cell-transcriptomics-reveal-a-hyperacute-cytokine-and-immune-checkpoint-axis-in-patients-with-poor-neurological-outcomes-after-cardiac-arrest~204, https://tinyurl.com/54t86fuu. Unsupervised clustering of the global single cell transcriptomic dataset revealed 6 major PBMC lineages: B cells, T cells, NK cells, monocytes, dendritic cells, and platelet-monocyte aggregates (**Figure 1B**), which were annotated by cell-type defining genes (Kunicki and Nugent, 2010; Schelker et al., 2017; Villani et al., 2017) (**Figure S2**). In t-SNE plots, patients with good outcomes co-clustered with cells from healthy subjects. At 6h post- OHCA, PBMC from patients with poor outcomes clustered separately and had a prominent loss of T cells (**Figure 1C-D**). This distinction between good and poor outcome patients was diminished by 48h post-CA. Principal component analysis (PCA) validated these trends at the per-patient level for NK cells and monocytes (**Figure 1E**). At 6h post-CA, PC1 alone largely separated patient sub-cohorts based on neurological outcome and healthy subjects. At 48h, both PC1 (36-40% of variance) and PC2 (12-18%) were required to segregate the patient sub-cohorts. Compared with healthy subjects, significant changes in the relative abundance of several cell type were observed early after CA (**Figure 1D, Figure S2**). Among the six major cell types, T cells had the greatest decrease in fractional abundance between control subjects (median [range]: 50 [45-65]%) and patients at 6h (good outcomes: 12 [6-44]%, P = 0.004; poor outcomes: 8 [4-29]%, P < 0.001).

**Figure 1.**
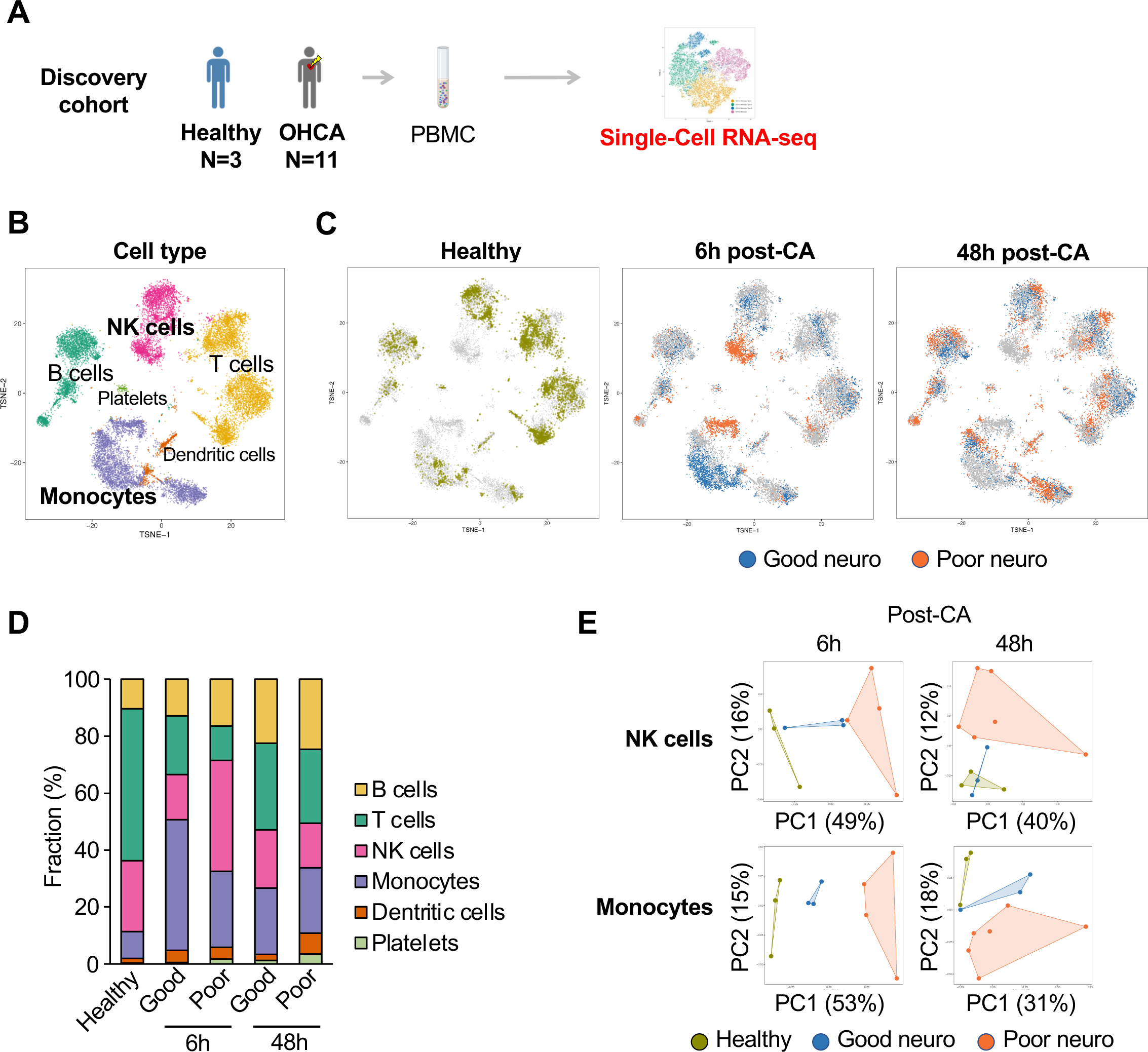
Distinct single-cell transcriptomic profiles distinguish innate immune cells from post-CA patients with poor or good neurological outcome early after CA. A) Approach for single-cell RNA-seq analysis of PBMC from patients after OHCA. Gross clustering of scRNA-seq dataset shown in t-SNE plots by B) cell type, and C) healthy subjects or time post-arrest for CA patients. D) Fraction of each cell type in PBMC. Good and Poor denotes good and poor neurological outcomes at hospital discharge, respectively. E) Principal component analysis of NK cells and monocytes at 6h and 48h post-CA, with individual patients and their subcohorts shown. CA, cardiac arrest; neuro, neurological outcomes; NK, natural killer; OHCA, out-of-hospital CA; PBMC, peripheral blood mononuclear cells; PC, principle component; RNA-seq, RNA sequencing.

Conversely, significant expansions of monocytes were observed early after CA compared to healthy subjects (healthy: 11 [5-12]%; good outcomes: 23 [10-51]%, P = 0.03; poor outcomes: 60 [9-69]%, P < 0.001). Other cell types did not show significant changes in their relative abundance. When fractional abundance was compared between neurological outcomes at 6h post-CA, there were no no significant difference in monocytes (P = 0.36), NK cells (P = 0.22), or T cells (P = 0.32).

### A monocyte subpopulation with increased expression of *NECTIN2* emerged by 6h post-arrest and associated with poor neurological outcome at hospital discharge

We first focused on monocytes, whose fractional abundance was significantly increased after CA compared to healthy subjects. Examining only monocytes in the scRNA-seq dataset, we applied the LCA clustering algorithm (Cheng et al., 2019) and identified four monocyte subclusters (**Figure 2A-B**). At 6h post-CA, monocyte cluster 4 emerged as the dominant cluster (median [range], 55 [13-81]% of monocytes) among patients that went on to have poor neurological outcomes. Monocyte cluster 4 was nearly absent from patients with good outcomes or healthy subjects (1.3 [0.3-2.3]% or 0.1 [0-0.4]% of monocytes, respectively, P < 0.001 compared to patients with poor outcomes). In contrast, monocyte cluster 2 was the dominant cluster in healthy subjects (74 [52-84]% of monocytes) and remained dominant at 6h post-arrest in patients that went on to have good neurological outcomes (54 [51-80]% of monocytes) but largely disappeared in patients with poor outcomes (5.5 [0-16]% of monocytes, P = 0.002 vs patients with good outcomes) (**Figure 2B)**. The difference in monocyte cell subpopulations between patients with poor and good outcomes was largely lost by 48h post-CA. At 48h post-CA, the majority of monocytes grouped in clusters 1 and 3 regardless of eventual neurological outcome. Interestingly, these monocyte clusters were not differentiated by well-established functional categories like “M1” and “M2” (Sica and Mantovani, 2012) (**Figure 2C**).

**Figure 2.**
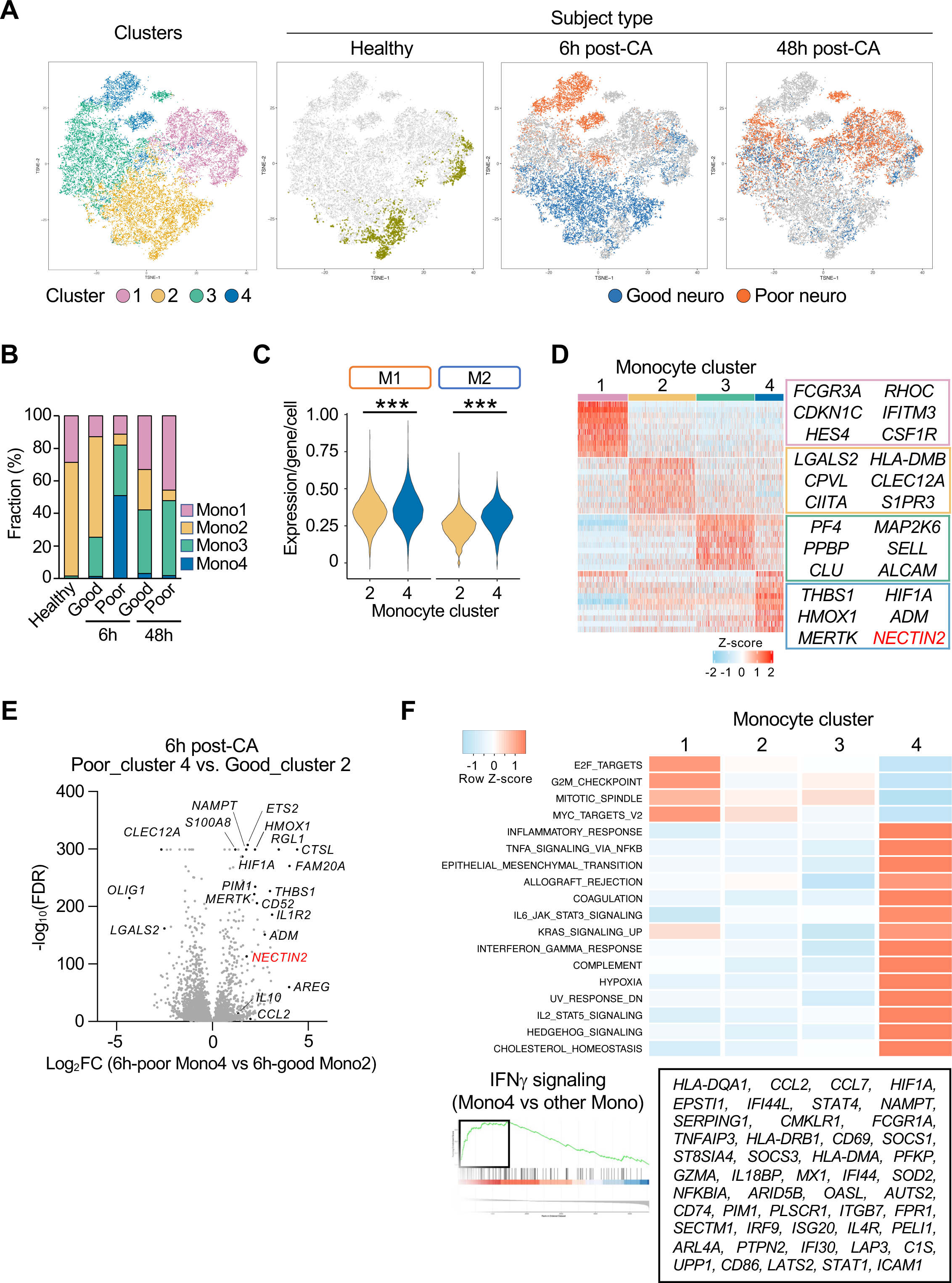
Hyperacute expansion of *NECTIN2*^+^ monocytes enriched for IFNγ-response pathways correlated with poor neurological outcomes after clinical CA. A) t-SNE plots of fine clustering of monocytes in the scRNA-seq dataset of PBMC. B) Proportions of monocyte subclusters. C) M1 and M2 gene signature scores for monocyte fine clusters 2 and 4. Average expression level for genes in the M1 or M2 signature (listed in Table S2) are shown. D) Cluster-defining genes for monocyte subclusters, with full list in Table S3. E) At 6h post-CA, differential gene expression analysis between the dominant monocyte clusters in patients with poor outcomes (cluster 4) and patients with good outcomes (cluster 2). See also Table S4. D-E) The Immune checkpoint gene confirmed in validation cohort is highlighted (red). F) Gene set enrichment analysis of monocyte clusters. Enrichment plot for IFNγ signaling shown. Z score was calculated by average gene expression of genes in the pathway from each cluster. See also Figure S3. CA, cardiac arrest; FC, fold-change; FDR, false discovery rate; neuro, neurological outcomes. C, Mann-Whitney U test, ***P < 0.001.

We examined the cluster-defining genes for these 4 monocyte subpopulations (**Figure 2D**, **Table S3**). Monocyte cluster 4 that enriched in patients with eventual poor outcomes at 6h post-CA overexpressed proinflammatory genes (e.g., *EDNRB*, *THBS1*, *TREM1*), anti-inflammatory genes (e.g., *HMOX1*), scavenger receptor genes associated with anti-inflammatory axes in some contexts (*CD163*, *MARCO, MERTK*) (Cai et al., 2018; Ghosh et al., 2011; Yang et al., 2016)), genes involved in invasion/migration (*TIMP1*), hypoxia-inducible genes (*HIF1A*, *ADM*, *HMOX1*), and inhibitory immune checkpoint molecule (*NECTIN2*). On the other side, genes defining cluster 2 (associated with healthy control and good outcome patients at 6h) are related to inflammation (*LGAL2*), antigen presentation (*CIITA*, *HLA-DMB*, *CPVL*), and genes with multiple immune and non-immune functions such as *CLEC12A* and *S1PR3* (Awojoodu et al., 2013; Neumann et al., 2014). *FCGR3A* (CD16)*^+^* non-classical monocytes (**Figure 2D**) are enriched in cluster 1 with similar abundance among patients with good or poor outcomes (**Figure 2B**, **Figure S3**). Most monocytes in cluster 3 were induced after 48h post-CA with similar fractions in good and poor outcome patients. It was defined in part by genes related to inflammation (*PF4* [CXCL4], *PPBP* [CXCL7]), cell death (*CLU*, *MAP2K6*), and adhesion (*SELL*, *ALCAM*).

To further delineate the key differences between cluster 2 (enriched in healthy and patients with good outcomes) and cluster 4 monocytes (enriched in patients with poor outcomes), we applied the NBID algorithm (Chen et al., 2018) that was designed for differential analysis in scRNA-seq after modeling the potential patient-specific effects (Chen et al., 2020) (**Figure 2E**, **Table S4**). Increased expression of prominent monocyte genes related to inflammatory (*THBS1*, *S100A8*, *S100A12*, *PIM1*, *PIM3*), immune checkpoint (*NECTIN2*), anti-inflammatory (*HMOX1*, *MERTK*, *IL1R2*, *CLEC12A*), and hypoxia-associated (*HIF1A*, *ADM*, *NAMPT*) genes were upregulated in monocyte cluster 4 associated with poor clinical outcomes. Gene set enrichment analysis (GSEA) on cluster-defining genes highlighted gene sets related to proinflammatory cytokines (*IFNG*, *TNFA*, *IL6*) and the hypoxia response in cluster 4 (**Figure 2F**).

In summary, monocytes in patients with poor neurological outcomes are dominated by a subpopulation characterized by increased expression of both pro- and anti-inflammatory genes, IFNγ-response genes, and the inhibitory immune checkpoint receptor Nectin-2. Although this monocyte subpopulation emerges and is enriched as early as 6h post-CA, a stage where the eventual neurological outcomes can remain unclear, it nevertheless strongly associates with poor neurological outcomes at hospital discharge. Since this monocyte signature was attenuated by 48h post-CA, single-cell analysis of monocytes emphasized the importance of hyperacute events occurring prior to 6h post-CA.

### A NK cell subpopulation with increased expression of *HAVCR2* and *TIGIT* emerged within 6h post-CA and associated with poor neurological outcomes

We next examined the NK cells, which were, along with monocytes, one of the two most abundant cell types at 6h after CA in patients with poor outcome at hospital discharge (**Figure 1D**). When NK cells are clustered separately, three subpopulations were evident (**Figure 3A**). Among them, NK cell cluster 1 dominated in patients with poor outcome at 6h post-CA (89.5 [85-95]%, **Figure 3B**), whereas cluster 2 was the major NK cell subpopulation in healthy subjects (85 [75- 94]%) and patients with good outcomes (55 [32-92]%). A third subpopulation was found at similar proportions among all subcohorts (4-19%) except at 6h post-CA in patients with poor outcomes (2 [1-6]%). Similar to monocytes, the NK cell profiles for CA patients with good outcomes and the profiles for CA patients with poor outcomes became more similar over time; at 48h post-CA, the distribution of NK cell subpopulations had trended back towards that of healthy subjects (**Figure 3B**).

**Figure 3.**
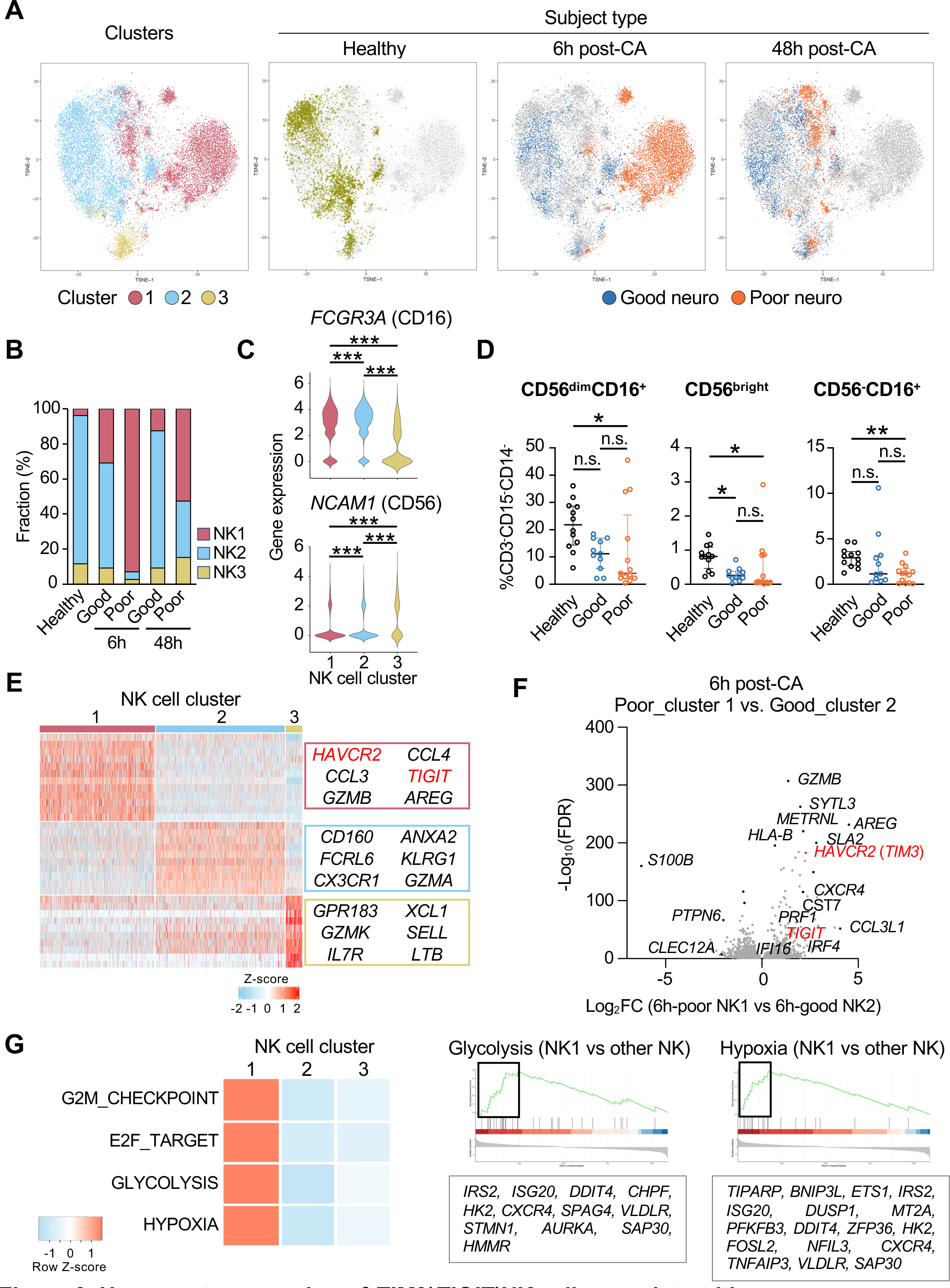
Hyperacute expansion of *TIM3^+^TIGIT*^+^NK cells correlate with poor neurological outcomes after clinical CA. **A)** t-SNE plots of fine clustering of NK cells in the scRNA-seq dataset of PBMC. **B)** Proportions of NK cell subclusters. **C)** Violin plots for *FCGR3A* (CD16) and *NCAM1* (CD56) expression in NK cell fine clusters. **D)** Quantification of NK cell subsets of healthy subjects and patients 6h post-CA by CD56 and CD16. Flow cytometry gating and 48h post-CA shown in Figure S4. **E)** Cluster-defining genes for NK cell subclusters, with full list in Table S5. **F)** At 6h post-CA, differential expression analysis between the dominant NK cell clusters in patients with poor outcomes (cluster 1) and in patients with good outcomes (cluster 2). See Figure S6. D-E) Immune checkpoint genes confirmed in validation cohort are highlighed (red). **G)** Gene set enrichment analysis of NK cell clusters. Z score was calculated by average gene expression of genes in the pathway from each cluster. Enrichment plots for glycolysis and hypoxia shown. CA, cardiac arrest; FC, fold-change; FDR, false discovery rate; neuro, neurological outcomes. C, D, Kruskal-Wallis test with Dunn’s multiple comparisons, *P < 0.05, **P < 0.01, ***P < 0.001.

Classical subset markers did not characterize the dominant NK cell clusters that differentiated patients by neurological outcome. *NCAM1* (CD56) and *FCGR3A* (CD16) were expressed at similar levels by NK cell clusters 1 and 2 (**Figure 3C**). Consequently, flow cytometry of patient PBMC showed similar fractions of CD56^bright/dim^ and CD16^+/-^ NK cell subpopulations between CA patients with good and poor outcome (**Figures 3D, S4**). Thus, we turned to cluster-defining genes to characterize these NK cell subpopulations. Genes defining NK cell cluster 1 (associated with poor outcome post-CA) include those associated with type 2 immune responses (*AREG*), inflammation (*CCL3*, *CCL4*, *IFNGR1*), cytotoxicity (*GZMB*) (Zaiss et al., 2015). Cluster 1 also upregulated two inhibitory receptors, the immune checkpoints *HAVCR2* [Tim-3] and *TIGIT* (Wolf et al., 2020) (**Figure 3E**, additional genes in **Table S5**). Cluster 2 (associated with good outcome post-CA) included genes associated with NK effector functions like IFNγ production and cytotoxicity (*CD160* (Tu et al., 2015), *FCRL6*, *GZMA* (Lieberman, 2010; Wang et al., 2013; Wilson et al., 2007). Cluster 2 also upregulated *KLRG1*, a negative regulator of NK cells (Lieberman, 2010; Wang et al., 2013) (**Figure 3E, Table S5**). Cells in cluster 3 had increased expression of genes characteristic of CD56^bright^ NK cells (*NCAM1, GPR183*, *GZMK*, *IL7R*, *XCL1*, *SELL*, *LTB*, and *KLRC1*). CD56^bright^ NK cells typically have increased production of cytokines and immunoregulatory functions, along with diminished cytotoxic activity(Smith et al., 2020). We employed DE and GSEA to compare the dominant NK cell clusters in patients with poor (cluster 1) and good outcomes (cluster 2). At 6h post-CA, differential expression (DE) analysis identified in cluster 1 additional genes in the same functional categories as its cluster-defining genes: inflammation (*CCL3L1*, *CCL4L2*, *CXCR4*, the IFNγ inducible genes *IFI16*, *IRF4*), cytotoxicity (*GZMB*, *PRF1*, *CST7*), and inhibitory receptors (*KIR2DL4*, *KIR* family, and the immune checkpoint receptors *HAVCR2* [Tim-3] and *TIGIT*) (Smith et al., 2020) (**Figure 3F, Table S6**). GSEA identified upregulation of genes associated with hypoxia and glycolysis (**Figure 3F**).

In summary, our scRNA-seq analysis revealed that, by 6h post-CA, a NK subpopulation dominated over 90% of NK cells in patients with eventual poor neurological outcomes. This subpopulation features a dysregulated profile of gene programs of inflammation, cytotoxicity, and inhibitory receptors (including the immune checkpoint receptors *HAVCR2* [Tim-3] and *TIGIT*), with a metabolic shift towards glycolysis (Wolf et al., 2020).

### Validation cohort: Nectin-2^+^ monocyte and Tim-3^+^ NK cell states are early molecular signatures of patients with poor neurological outcomes after CA

To validate our findings in scRNA-seq in the discovery cohort, we enrolled a second cohort of patients with OHCA (N = 28) (**Table S7**). We tested the hypothesis that Nectin-2^+^ monocytes and Tim-3^+^ NK cells are early molecular signatures of eventual poor neurological outcomes after CA. Nectin-2 was a cluster-defining gene for monocyte cluster 4, the dominant monocyte subpopulation found in patients with poor neurological outcomes (**Figure 2B**). Nectin-2 is a ligand of the inhibitory, immune checkpoint receptor TIGIT (Tahara-Hanaoka et al., 2006). TIGIT was a cluster-defining gene for NK cell cluster 1, the dominant NK cell subpopulation in CA patients with poor outcomes (**Figure 3E**). We sorted innate immune cells from post-CA patients by the immune checkpoint receptors Nectin-2 (for monocytes) and Tim-3 (for NK cells) and performed bulk RNA-seq (**Figure S5**). Flow cytometry analysis confirmed that the proportion of Nectin-2^+^ monocytes was significantly higher in patients with poor outcomes (49% [14-57] of monocytes, n = 10) at 6h post-CA, compared to both patients with good outcomes (32% [4-41], n = 10, P = 0.03) and healthy subjects (17% [4-39], N = 11, P = 0.002) (**Figure 4B**). In line with scRNA-seq findings, patients with good or poor outcomes became similar over time, as both outcomes had similar proportions of Nectin-2^+^ monocytes by 48h post-CA (P = 0.90). At 6h post-CA, significantly more Tim-3^+^ NK cells were recovered from patients with poor outcomes (8% [3-82] of NK cells, n = 13) than both patients with good outcomes (0.9% [0.4-5], n = 11, P < 0.001) and healthy subjects (1% [0.4-6], N = 12, P = 0.002) (**Figure 4C**). At 48h post-CA, a similar trend was observed (P = 0.04).

**Figure 4.**
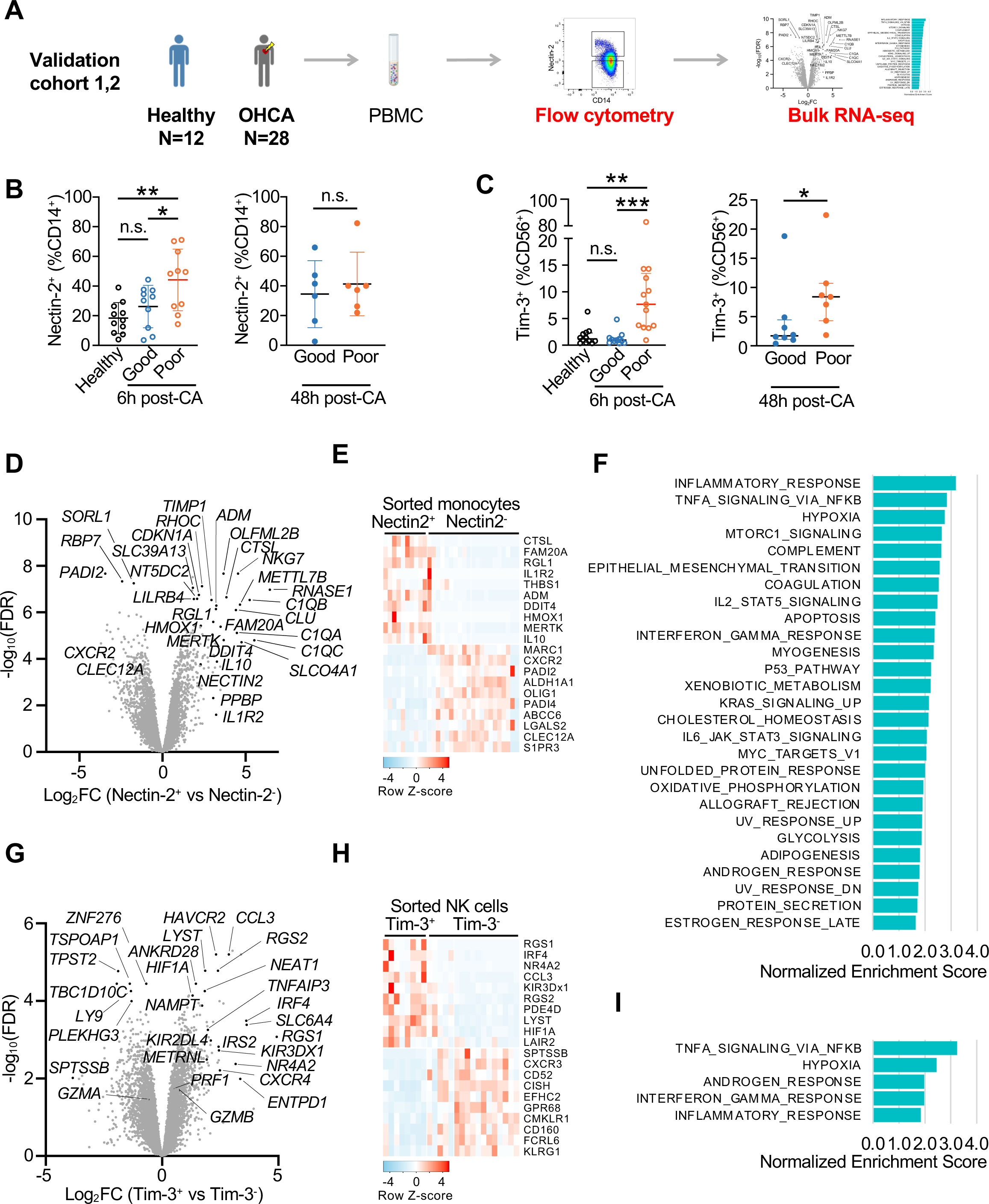
Validation cohort confirms expansion of Nectin-2^+^ monocyte and Tim-3^+^NK cell states in CA patients with poor neurological outcomes. A) Approach for confirmation of scRNA-seq results in validation cohort of CA patients, with clinical characteristics in Table S7. B) Flow cytometry analysis of Nectin-2^+^ monocytes. Per patient % Nectin-2^+^ monocytes are shown. **C)** Flow cytometry analysis of Tim-3^+^ NK cells. Per patient % Tim-3^+^ NK cells are shown. B-C) Flow cytometry gating strategy shown in Figure S5. **D)** Nectin-2^+^ and Nectin-2^-^ CD14^+^ monocytes were sorted by flow cytometry and assessed by bulk RNA-seq. Differential expression (DE) of genes were calculated for Nectin-2^+^ monocytes from patients with eventual poor outcomes compared to Nectin-2^-^ monocytes from patients with eventual good outcomes (measured at 6h post-CA) or healthy subjects. Fold-change (FC) is shown for patients at 6h-post CA or healthy subjects. See also Table S8. **E)** Heatmap of bulk RNA-seq dataset is shown for sorted Nectin-2^+^ monocytes from 6h poor outcomes vs Nectin-2^-^ monocytes from 6h good outcomes and healthy subjects. Genes shown are differentially expressed genes identified in the scRNA-seq dataset between Nectin-2^+^ monocyte cluster 4 and Nectin-2^-^ monocyte cluster 2. See also Figure S5. **F)** Gene set enrichment analysis (GSEA) of the bulk RNA-seq of sorted Nectin-2+/- monocytes using the same comparison as in (4D). **G)** Tim-3^+^ and Tim-3^-^ NK cells were sorted by flow cytometry and assessed by bulk RNA-seq. Differential expression (DE) of genes were calculated for Tim-3^+^ NK cells from patients with poor outcomes post-CA compared to Tim-3^-^ NK cells from patients with good outcomes post-CA or healthy subjects. FC is shown for patients at 6h post-CA or healthy subjects. See also Table S9. **H)** Heatmap of bulk RNA-seq dataset is shown for sorted Tim-3^+^ NK cells from 6h poor outcomes vs Tim-3^-^ NK cells from 6h good outcomes and healthy subjects. Genes shown are differentially expressed genes identified in the scRNA-seq dataset between Tim-3^+^ NK cluster 1 and Tim-3^-^ NK cell cluster 2. See also Figure S5. **I)** GSEA of the bulk RNA-seq of sorted Tim-3+/- NK cells using the same comparison as in 4G. CA, cardiac arrest; FC, fold-change; FDR, false discovery rate; OHCA, out-of-hospital cardiac arrest; RNA-seq, RNA sequencing. Good and Poor denote good and poor neurological outcomes at hospital discharge, respectively. B, C, Kruskal-Wallis test with Dunn’s multiple comparisons or Mann-Whitney U test, *P < 0.05, **P < 0.01, ***P < 0.001.

We next assessed whether Nectin-2^+^ monocytes faithfully recaptured the molecular signatures of the dominant monocyte subpopulation (cluster 4) defined by scRNA-seq. DE analysis using bulk RNA-seq profiling of the sorted Nectin-2^+^ and Nectin-2^-^subpopulations at 6h post-CA replicated the transcriptomic difference between the cluster 4 monocytes (enriched in patients with poor outcomes) and the cluster 2 monocytes (enriched in healthy subjects and patients with good outcomes) in the scRNA-seq DE analysis. As examples, both monocyte cluster 4 (scRNA-seq, discovery cohort) and flow cytometry sorted Nectin-2^+^ monocytes (bulk RNA-seq, validation cohort) had increased expression of *NECTIN2*, *IL10, MERTK*, *THBS1*, and *HMOX1,* and down-regulation of *CLEC12A*, *HLA-DMB*, and *LGAL2* (**Figure 4D**, **Table S8**). Genes differentially expressed among the dominant monocyte clusters of poor (Nectin-2^+^) and good outcomes (Nectin-2^-^) in scRNA-seq dataset were also differentially expressed in bulk RNA-seq dataset of sorted Nectin-2^+^ and Nectin- 2^-^ monocytes in bulk RNA-seq dataset (**Figure 4E**). Moreover, across all genes, the fold change in the DE analysis showed a high correlation between scRNA-seq in the discovery cohort and bulk RNA-seq in the independent validation cohort, with ρ = 0.82 [0.77-0.85] (P < 0.001) (**Figure S5**). In bulk RNA-seq of Nectin-2^+^ monocytes, GSEA identified upregulation of pathways involving inflammation (including TNFα and IFNγ responses), hypoxia, and metabolic upregulation (of both oxidative phosphorylation and glycolysis) (**Figure 4F**). These findings suggest a dysregulated inflammatory response and metabolic shift within Nectin-2^+^ monocytes, a finding that correlates closely with those found in the scRNA-seq analysis (**Figure 2E-F**). This result suggests that Nectin-2 alone was a sufficient marker to identify the dominant monocyte subpopulation in patients with poor neurological outcomes at hospital discharge.

Similarly, Tim-3-based cell sorting and bulk RNA-seq profiling demonstrated that Tim-3 is a robust marker to identify the cluster 1 NK cells that are predominant in PBMC collected at 6h post-CA from patients with poor outcomes (**Figure 4G**). Overall, the transcriptomic difference between Tim-3^+^ and Tim-3^-^ NK cells in the validation cohort accurately captured the difference between cluster 1 and cluster 2 NK cells from scRNA-seq of the discovery cohort (ρ = 0.83 [0.79-0.87], P < 0.001), Spearman’s correlation, **Figure 4H**, **Figure S5**). As in the discovery cohort, GSEA of Tim- 3^+^ NK cells from the validation cohort revealed enrichment in gene sets encoding inflammatory responses (including TNFα and IFNγ responses) and hypoxia (**Figure 4I**).

In summary, single-cell approaches identified two novel innate immune cell subpopulations that emerged within 6 hours after CA and were associated with subsequent neurological outcomes at hospital discharge. The enrichment of Nectin-2^+^ monocyte and Tim-3^+^ NK cells in patients with poor outcomes was further confirmed in a validation patient cohort at the protein level and by bulk RNA-seq profiling. GSEA in the validation cohort confirmed the scRNA-seq finding that IFNγ and TNFα inflammatory axes are major pathways in these innate immune cell subpopulations.

### Interactome analysis identifies candidates for monocyte and NK cell crosstalk

Monocytes and NK cells modulate the early immune response by either cell to cell contact or release of soluble factors in both an autocrine and paracrine fashion (Kossmann et al., 2013; Welte et al., 2006). Using interactome analysis by the NicheNet algorithm (Browaeys et al., 2020), we explored ligand-receptor axes that may mediate crosstalk between these two cell types after CA. The analysis of “monocyte ligand-NK cell receptor” pairs highlighted opposing proinflammatory and anti-inflammatory axes. NicheNet identified monocyte production of inflammatory cytokines (e.g., *IL15*, *IL18*) and *TNFSF*s (*TNFSF10* [TRAIL], *TNFSF12* [TWEAK], *TNFSF13B* [BAFF], and *TNFSF14* [LIGHT]), and chemokines (*CCL2*, *CCL5*, *CCL7*) that could drive activation, migration, and metabolic shifts in NK cells (Fan et al., 2006; Keppel et al., 2015; Marçais et al., 2014; Robertson, 2002; Shan et al., 2006; Smyth et al., 2001) (**Figure 5A**). The converse analysis of “NK cell ligand-monocyte receptor” pairs highlighted NK cell expression of *IFNG*, along with other cytokines known to activate or attract monocytes (**Figure 5B**). Like DE and GSEA, interactome analysis featured *IFNG*-related pathways. For example, IL-15 and IL-18 are known to drive production of IFNγ by NK cells (Fehniger et al., 1999; Fleetwood et al., 2007), and IFNγ can induce expression of TWEAK by monocytes, which in turn can repress production of IFNγ by NK cells (Maecker et al., 2005; Nakayama et al., 2000). We also identified anti-inflammatory axes enriched in CA patients with poor outcome: the inhibitory immune checkpoint axis *NECTIN2*-*TIGIT*, *IL10*, and *TGFB1*. The analysis also identified potential interaction between Nectin-2 and CD226, which can activate cytotoxic lymphoid cells (Gilfillan et al., 2008) and interact with the Nectin-2/TIGIT axis (Johnston et al., 2014). In interactome analysis for autocrine loops in monocytes (**Figure 5C**), we re-identified several axes found between NK cells and monocytes, including *IL10*. Together, the interactome analysis called attention to the early emergence of both proinflammatory (particularly IFNγ) and anti-inflammatory axes (IL-10 and immune checkpoint receptors) between NK cells and monocytes in patients with poor neurological outcomes after CA.

**Figure 5.**
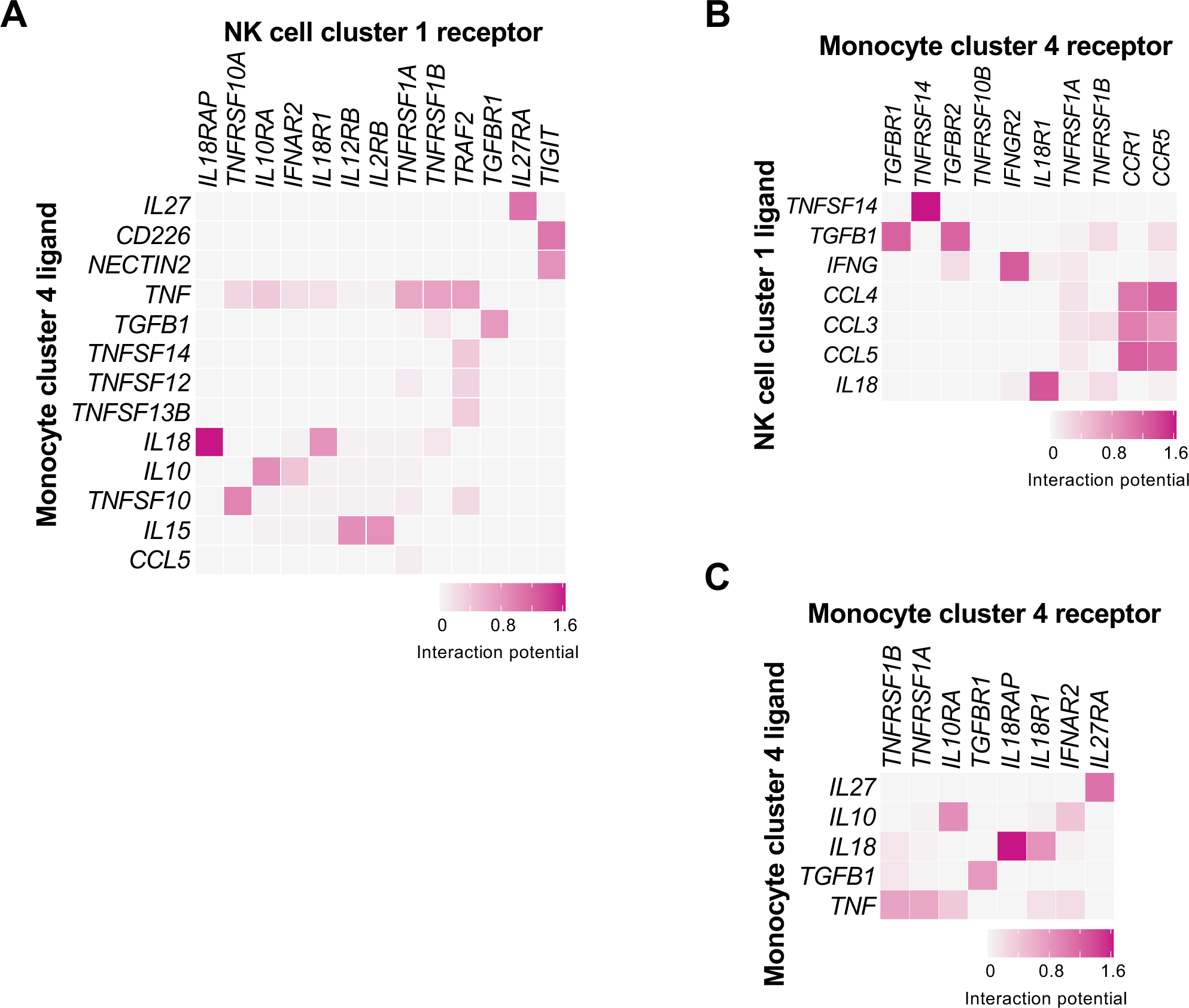
Interactome analysis of *NECTIN2^+^* monocyte and *TIM3^+^TIGIT^+^*NK cell states identifies immune checkpoint and pro- and anti-inflammatory axes in cardiac arrest patients with poor neurological outcomes. The NicheNet algorithm was applied to the scRNA-seq dataset and, at 6h post-cardiac arrest, CA patients with poor neurological outcome are compared to patients with good neurological outcome. Significantly increased interactions are shown for: A) Ligands secreted by *NECTIN2^+^*monocytes binding to receptors on *HAVCR2^+^TIGIT^+^* NK cells. NK cells ligands to receptors on monocytes. B) Ligands secreted by *HAVCR2^+^TIGIT^+^*NK cells binding to receptors on *NECTIN2^+^* monocytes. C) Ligands secreted by *NECTIN2^+^* monocytes binding to receptors on *NECTIN2^+^* monocytes (autocrine).

### A hyperacute (6h post-arrest) increase in plasma cytokine and chemokine levels associated with poor neurological outcomes after OHCA

Next, we selectively validated clinically relevant predictions of the interactome analysis. To increase rigor, we enrolled a third cohort of cardiac arrest patients to measure predicted cytokines and chemokines. Plasma was assessed by multiplex ELISA in a total of 47 post-CA patients (18 patients with good and 29 with poor outcomes) and healthy controls (n = 15) (**Figure 6A, Table S10**). In line with the interactome analysis, plasma levels of IFNγ, TNFα, IL-10, IL-18, and CCL2 were significantly increased in CA patients with poor outcome at 6h after CA (**Figure 6B**). Similar to our scRNA-seq findings, the differentiation in cytokine/chemokine levels between patients with poor and good outcomes was largely lost by 48h after CA. We also measured selected cytokines upstream and downstream of IFNγ. Like IL-18 and IL-12 can stimulate IFNγ production by lymphoid cells. IL-12p40 distinguished patients by neurological outcome at 6h but not 48h post-arrest. However, IL-12p70 was comparable across groups and timepoints. IFNγ can induce release of IL-1β (Masters et al., 2010), which has been associated with poor outcomes after CA in a previous study (Asmussen et al., 2021). At 6h post-CA, plasma IL-1β showed a trend toward increase among patients with poor outcomes (P = 0.07) and became comparable across groups at 48h post-CA. The *NECTIN2^+^* cluster (scRNA-seq) and flow cytometry sorted Nectin-2^+^ monocytes (bulk RNA-seq) identified an IL-6 response at 6h post-CA in patients with poor outcome. Indeed, plasma IL-6 levels did distinguish patients by neurological outcome at 6h but not 48h post-arrest (**Figure 6B)**. Notably, the cytokines and chemokines identified by our single-cell and bulk RNA-seq analyses (e.g., IFNγ, TNFα, IL-10, IL-18) showed more significant and striking discrimination between patients with good and poor neuro outcome than cytokines that were not identified by RNA-seq but tested because they were reported in the literature (i.e., IL-1β, IL-12 isoforms).

**Figure 6.**
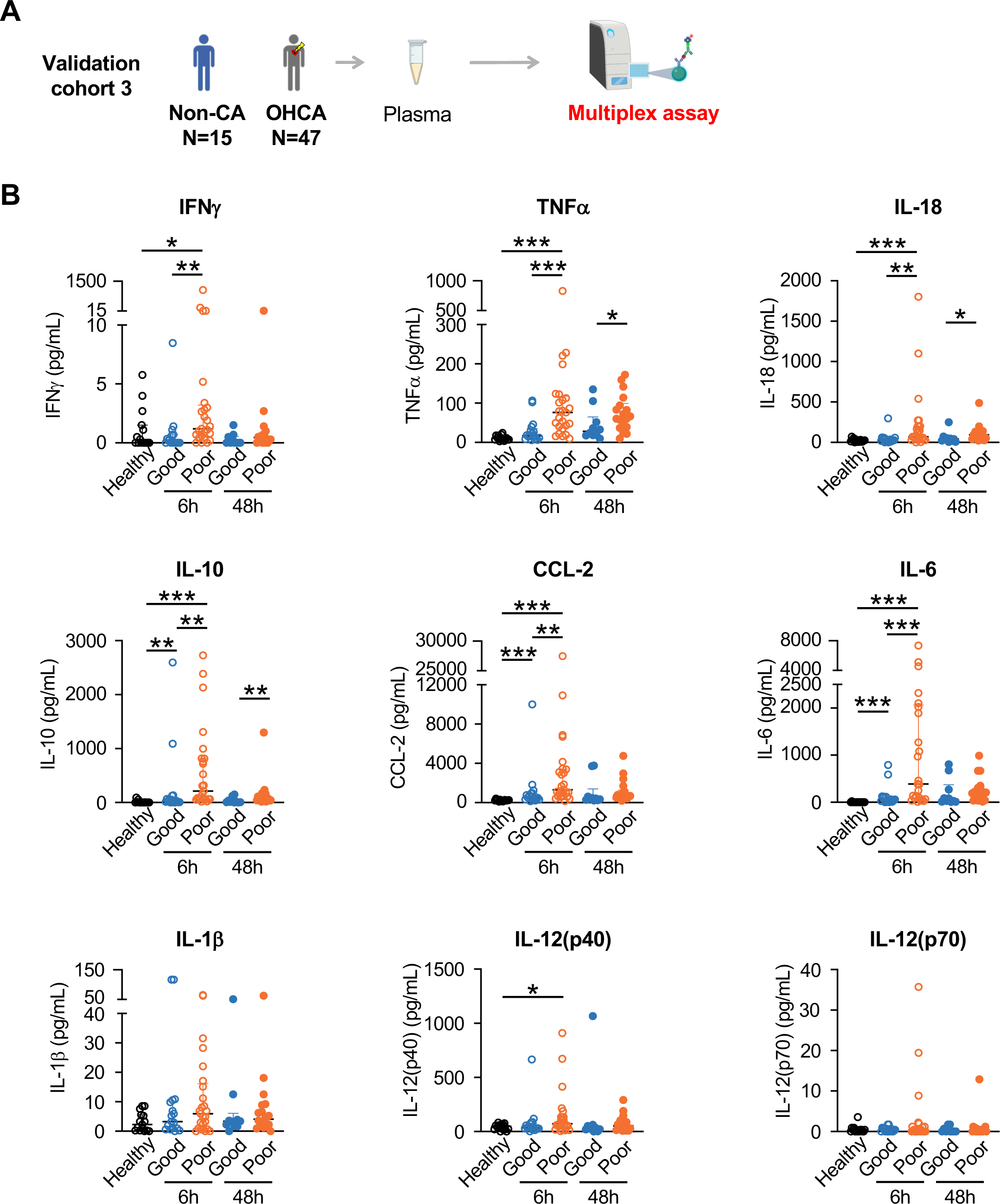
Hyperacute elevation in plasma levels of pro- and anti-inflammatory cytokines distinguish CA patients with poor neurological outcomes. A) Approach to assessment of cytokines identified by interactome analysis in a separate validation cohort of CA patients, with clinical characteristics in Table S10. B) Cytokines and chemokines measured by multiplex ELISA of plasma from CA patients with good or poor neurological outcome at time points post-CA or healthy subjects. CCL, C-C Motif Chemokine Ligand; IFNγ, interferon-gamma; IL, interleukin; TNFα, tumor necrosis factor-alpha. Mann-Whitney U test with Benjamini-Hochberg correction for multiple comparisons, *P < 0.05, **P < 0.01, ***P < 0.001. Good and Poor denotes good and poor neurological outcomes, respectively.

### In cardiac arrest, monocyte-NK cell crosstalk is governed by IFNγ, IL-10, and Nectin-2

Our complementary approaches in three independent patient cohorts repeatedly demonstrated that patients with poor neurological outcomes after cardiac arrest had very early elevation of immune checkpoint receptors and pro- and anti-inflammatory cytokines (**Figures 4, 6**). We next examined in functional studies if the immune checkpoint and cytokine axes interacted (**Figure 7A**). First, we asked if these hyperacute cell subpopulations could be reproduced in vitro. We exposed healthy PBMC to the cytokine milieu identified by our interactome analysis. Prior studies explored the regulation of Nectin-2’s receptors (TIGIT, DNAM1, and CD112R); but the inducers of Nectin-2’s expression are not well-defined. Among cytokines identified by the scRNA-seq interactome analysis, IFNγ and IL-10 upregulated expression of *NECTIN2* on monocytes (**Figure 7B**). Inspired by the enrichment of hypoxia response genes in patients with poor outcomes after cardiac arrest (**Figure 2F, 4E)**, we also treated monocytes with molidustat, a prolyl hydroxylase inhibitor that stabilizes hypoxia-inducible factor (HIF) and is used as a “chemical model of hypoxia (Flamme et al., 2014)”. Molidustat had little impact on the expression of *NECTIN2* (**Figure 7B**), which suggested that HIF-dependent effects of hypoxia are not the direct driver of the *NECTIN2* expression after CA. Although IFNγ and IL-10 are considered pro- and anti-inflammatory cytokines with “opposing effects,” here, these cytokines had an additive effect on induction of *NECTIN2* expression. In contrast to their effect on expression of *NECTIN2*, the “cooperation” between IFNγ and IL-10 was not seen for pro- and anti-inflammatory cytokines, as expected (**Figure S5**). Next, we confirmed this finding at the protein level. Co-stimulation with IFNγ and IL-10 had an additive effect on increasing expression of Nectin-2^+^ on monocytes as measured by flow cytometry (**Figure 7C**).

**Figure 7.**
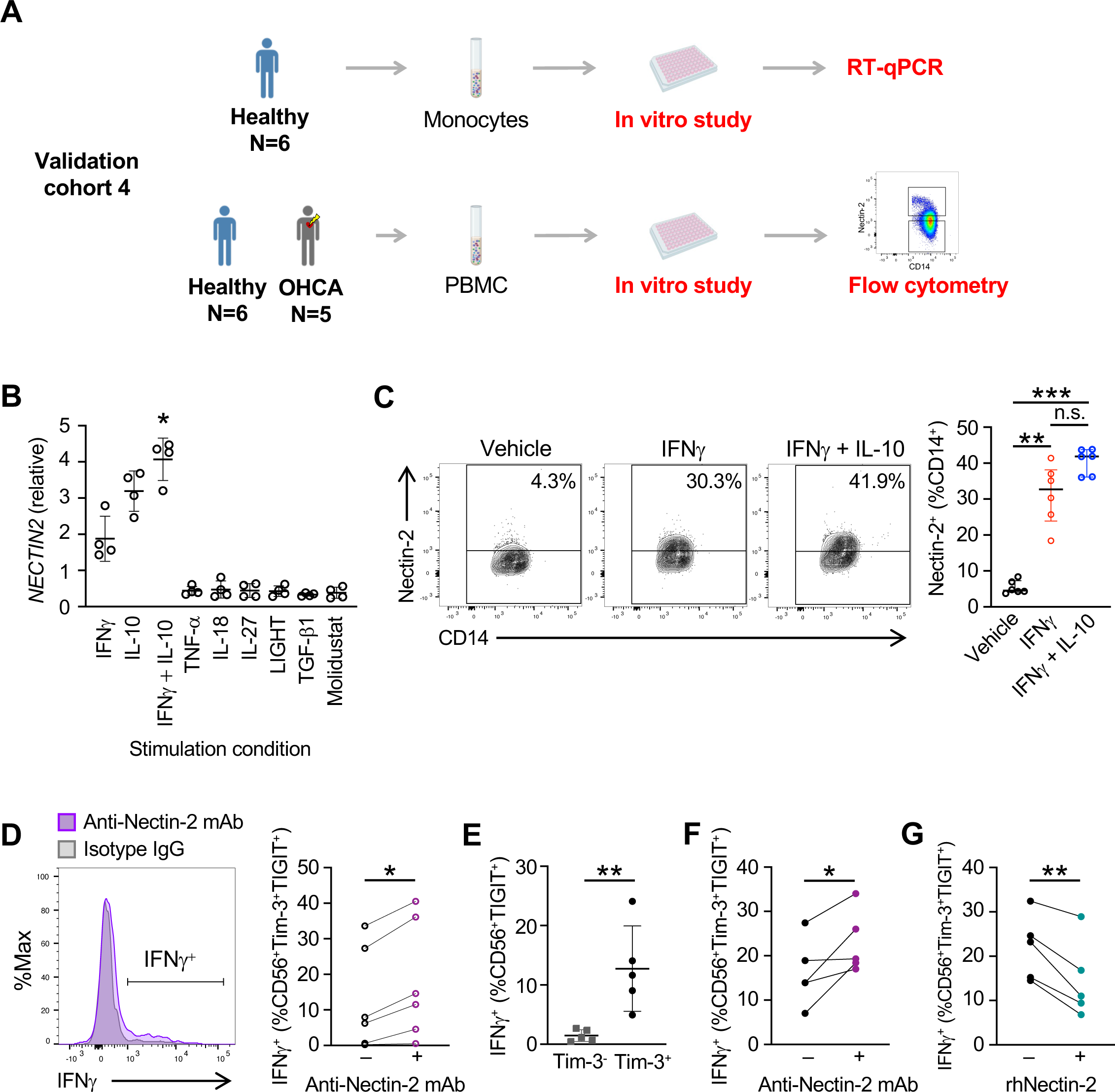
Nectin-2 mediates a negative feedback loop to limit production of IFNγ by Tim- 3^+^TIGIT^+^ NK cells. **A)** Approach for (7B) in vitro study of monocytes from healthy human subjects, (7C-D) in vitro study of PBMC from healthy subjects, and (7E-G) ex vivo study of PBMC from patients at 6h post-CA with poor neurological outcome. **B)** Monocytes from healthy subjects were treated in vitro with cytokines identified by interactome analysis (Figure 5). *NECTIN2* mRNA expression levels measured in monocytes by quantitative PCR (qPCR) are shown. Expression is normalized to housekeeping gene and to unstimulated control for each subject. **C)** Flow cytometric analysis of Nectin-2^+^ monocytes from healthy subjects after in vitro stimulation. Representative plots and % Nectin-2^+^ monocytes (per patient) are shown. **D)** Intracellular IFNγ in Tim-3^+^TIGIT^+^ NK cells measured by flow cytometry after PBMC from healthy subjects treated in vitro with cytokine (IFNγ+IL-10) stimulation after treatment with anti-Nectin-2 blocking monoclonal antibody (mAb) or isotype control. Representative plot and % IFNγ^+^Tim-3^+^TIGIT^+^ NK cells (per patient) shown. **E)** % IFNγ^+^Tim3^+^TIGIT^+^ and IFNγ^+^Tim-3^−^TIGIT^+^ NK cells at 6h post-CA in patients with poor neurological outcome, measured by flow cytometry. **F)** % IFNγ^+^Tim-3^+^TIGIT^+^ NK cells in the presence or absence of Anti-Nectin-2 mAb. **G)** Intracellular IFNγ levels of early post-CA Tim- 3^+^TIGIT^+^ NK cells after treatment with recombinant human (rh)-Nectin-2. OHCA, out-of-hospital cardiac arrest; B, C, Welch ANOVA test with Dunnett’s T3 multiple comparisons; D, F, G, ratio paired t-test; E, unpaired t-test. *P < 0.05, **P < 0.01, ***P < 0.001.

We next asked the converse question, whether immune checkpoint receptors modulate cytokines in CA. Nectin-2 binding to TIGIT inhibits inflammation or cytotoxicity by T cells in cancer (Ho et al., 2021; Lozano et al., 2020; Oshima et al., 2013) and inhibits IFNγ production by NK cells in viral hepatitis (Bi et al., 2014b, 2014a; Stanietsky et al., 2009); but the role of Nectin-2 has not been described in acute critical illness. We hypothesized that Nectin-2^+^ monocytes represented a negative feedback loop to promote the resolution of inflammation after cardiac arrest. As proof-of-principle, we stimulated PBMC from healthy subjects in vitro with IFNγ and IL-10 and demonstrated that the addition of anti-Nectin-2 blocking monoclonal antibody (mAb) increased IFNγ production by NK cells (**Figure 7D)**. We then tested this hypothesis in ex vivo studies of PBMC from post-CA patients. Our scRNA-seq and bulk RNA-seq results identified Tim-3^+^TIGIT^+^ NK cells as the source of *IFNG*. However, recent studies in other contexts have shown that Tim-3 expression can mark either NK cells that are potent producers of IFNγ (Gleason et al., 2012; Ndhlovu et al., 2012) or NK cells that resemble “exhausted” T cells (da Silva et al., 2014; Xu et al., 2015). First, we confirmed that Tim-3^+^TIGIT^+^, but not Tim-3^−^TIGIT^+^ NK cells were the dominant source of IFNγ production among NK cells in CA (**Figure 7E**). Addition of anti-Nectin-2 mAb increased the IFNγ production by Tim- 3^+^TIGIT^+^ NK cells in PBMC from post-CA patients (**Figure 7F**). In the opposite intervention, NK cells from CA patients produced less IFNγ when incubated with recombinant Nectin-2, an agonist of the receptors binding Nectin-2 (**Figure 7G**). In summary, IFNγ and IL-10 cooperatively induce expression of the immune checkpoint receptor Nectin-2 on monocytes. In turn, Nectin-2 down- regulates production of IFNγ by Tim-3^+^TIGIT^+^ NK cells after CA. These results demonstrate how a mixed pro- and anti-inflammatory cytokine milieu can trigger a mechanism promoting the resolution of inflammation after cardiac arrest (**Figure S1**).

## Discussion

In this study, we defined the immunological network of innate immune cells following clinical OHCA at single-cell resolution. In the hyperacute phase at 6h after CA, Nectin-2^+^ monocyte and Tim-3^+^TIGIT^+^ NK cell subpopulations were highly expanded among patients with poor neurological outcomes at hospital discharge. These innate immune cell states were defined de novo by scRNA-seq and did not fit classical schemes like M1/M2 monocytes. Notably, the hyperacute difference in innate immune endotypes between CA patients with poor and good neurological outcomes was largely attenuated by 48h post-arrest. Interactome analysis suggested that cytokines and immune checkpoints mediated crosstalk among innate immune cells. The findings from scRNA-seq were confirmed in two validation cohorts of OHCA patients at the protein level and at the global transcriptomic level by bulk RNA-seq of cell subpopulations. We then dissected the interaction between cytokine and immune checkpoint axes. The proinflammatory cytokine IFNγ and the anti-inflammatory cytokine IL-10 cooperatively induced expression of Nectin-2 in monocytes; in a negative feedback loop, Nectin-2 was a brake on IFNγ production by NK cells. In ex vivo studies of PBMC from CA patients, blockade of Nectin-2 increased IFNγ production by Tim-3^+^TIGIT^+^ NK cells. In proof-of-principle for a therapeutic intervention, treatment with recombinant Nectin-2-Fc fusion protein inhibited IFNγ production by Tim-3^+^TIGIT^+^ NK cells. In the limited studies of the immunology of clinical CA to date, the focus has been on potentially pathogenic, proinflammatory axes. Here, we highlight a mechanism for the resolution of inflammation, in which pro- and anti-inflammatory cytokines cooperatively induce a negative feedback loop.

In CA, the immune system has been targeted therapeutically in a limited fashion. Most clinical trials used untargeted treatments, such as corticosteroids and targeted temperature management, and have had mixed efficacy on outcomes (Donnino et al., 2016; Shah and Mitra, 2021). A recent clinical trial in cardiac arrest showed that the IL-6 receptor antagonist tocilizumab reduced plasma C reactive protein levels and myocardial injury, but tocilizumab did not improve survival or neurological outcomes (Meyer et al., 2021). We did identify an IL-6 response in monocytes associated with poor neurological outcome, but we also found a strong association between other inflammatory cytokines, such as IFNγ and TNFα, and neurological injury. Our results could motivate clinical trials targeting other cytokines like IFNγ. However, it is plausible that multiple cytokines contribute to CA, and, therefore, a need to concurrently intervene against several cytokines may limit the success of such an approach. Our scRNA-seq study suggests that a more effective strategy may be upstream intervention by targeting the cellular source of cytokines, such as Tim-3^+^TIGIT^+^ NK cells, and inhibiting multiple downstream effector molecules with a single treatment like recombinant Nectin-2.

Our study underscores the importance of timing in proposed therapeutic interventions in CA. In our study, hyperacute (<6h post-arrest) innate immune and cytokine responses were associated much more strongly with poor neurological outcomes than the immune profile at 48h post-arrest.

Although 48h would still be considered an early time point for critical illness, our study suggests that the very earliest innate immune response may have a profound influence in the pathogenesis of neurological injury and mortality. Thus, the time to intervention, measured in hours, might be critical for clinical trials. For example, in a major clinical trial comparing two target temperatures (Dankiewicz et al., 2021) the average time to reach the target temperature was eight hours post-CA, that is, after the immune endotypes from our study were established. However, unlike sepsis in which the infection may have begun days prior to hospital presentation, because of its discrete clinical onset, CA offers an opportunity to intervene just minutes after activation of the immune response.

Our study introduces immune checkpoints, such as Nectin-2, as a potential treatment class for very early intervention after CA. Also called CD112, Nectin-2 has described roles in cellular adhesion, proliferation, and differentiation (Takai and Nakanishi, 2003). The regulation of *NECTIN2* expression is not well-defined, so our finding that IFNγ and IL-10 cooperatively induce expression of *NECTIN2* may also inform other diseases. Nectin-2 has gained attention for its immunomodulatory properties, particularly in oncology (Ho et al., 2021; Lozano et al., 2020; Oshima et al., 2013). By binding the receptors TIGIT and CD112R, Nectin-2 can inhibit cytotoxic T cell activation and proliferation (Whelan et al., 2019; Zhu et al., 2016). While immune checkpoint pathways have been extensively studied among T cells, their function in innate immunity and contribution to disease pathology outside of cancer has only recently drawn attention. This study introduces a TIGIT-Nectin-2 axis for inhibition of NK cell function in critical illness. Like most translational studies of CA, our study is limited to peripheral circulation. Immune checkpoints have not been examined in experimental cardiac arrest, but experimental single-organ IRI models suggest that immune checkpoints may operate in multiple mechanisms in end organs during local ischemia. In experimental stroke, scRNA-seq did not highlight immune checkpoints; but the majority of monocytes infiltrating ischemic brain parenchyma did express *Ctsl and Thbs1*, two genes overexpressed in the monocyte cluster associated with poor outcomes in our study of CA (Zheng et al., 2022). In scRNA-seq analysis of human liver 2h after transplantation and reperfusion, a CD8^+^ T cell subset had enrichment of the PD-1 signaling gene set, but there were no immune checkpoints highlighted in other cell subpopulations (Wang et al., 2021). Blockade of the PD-1 /PD-L1/2 axis worsened tissue injury in both experimental cardiac and renal IRI, via macrophage polarization or regulatory T cell functions, respectively (Jaworska et al., 2015; Xia et al., 2020). These results suggest that incomplete control of inflammation by endogenous immune checkpoints is common to both global and single-organ IRI but raise the possibility that there are distinct mechanisms in global and single-organ IRI.

Our study highlights the hyperacute rise of IFNγ in CA patients with poor neurological outcomes. TNFα, IL-6, and IL-10, but not IFNγ, have been prominently associated with mortality in clinical CA (Adrie et al., 2002; Fries et al., 2009) The role of IFNγ may have been previously underappreciated in CA because its levels rapidly fall after the hyperacute window, whereas TNFα and IL-6 remain markedly elevated at 48h post-arrest (**Figure 6B**). Further, biologically significant plasma levels for IFNγ are often in a low range of absolute values. For example, in our prior study of patients in the first 48h of sepsis, the majority of plasma IFNγ values were in the 2-10 pg/mL range (as in this cardiac arrest study) despite the known robust IFNγ response in sepsis (Kim et al., 2020). Blockade of IFNγ has not been examined in experimental CA. However, NK cells and IFNγ drives injury in some single-organ IRI models, such as experimental stroke (Bajpai et al., 2019; Chu et al., 2015; Gan et al., 2014; Planas, 2018; Zhang et al., 2014) and renal IRI (Daemen et al., 1999; Day et al., 2006), although IFNγ blockade did not ameliorate experimental myocardial IRI (Homma et al., 2013). IL-10 is protective in experimental myocardial ischemia (Homma et al., 2013), stroke (de Bilbao et al., 2009; Tukhovskaya et al., 2014), and renal IRI (Tang et al., 2020; Yan et al., 2020). Studies of experimental IRI have not examined whether IL-10 amplified its effect via induction of immune checkpoints, as seen in our study. While mechanisms differ from our study of clinical CA, experimental IRI models have also identified both deleterious and protective innate immune subpopulations. For example, inflammatory Ly6c^hi^ monocytes infiltrate end-organs in response to CCL2 in myocardial and renal IRI (Bajpai et al., 2019; Furuichi et al., 2003; Hilgendorf et al., 2014; Song et al., 2018), but Ly6c^hi^ monocytes can also be protective in experimental stroke (Chu et al., 2015) or myocardial IRI (Hilgendorf et al., 2014).

Our study corroborated some findings from the limited immunophenotyping of prior studies in clinical CA (Asmussen et al., 2021; Kim et al., 2018; Miyatake et al., 2019; Ryzhov et al., 2019; Villois et al., 2017; Wei ser et al., 2017). For example, a prior study (Asmussen et al., 2021) associated reduced expression of HLA-DR on monocytes with poor outcomes after CA. This finding is in line with our scRNA-seq study, in which monocyte cluster 4 (the dominant cluster in poor outcome patients) had downregulation of HLA-DR genes. Our study may also provide insight into a prior study of whole blood bulk transcriptional profiling, which found upregulation of the inflammatory genes *CTSL*, *RGS1*, and *DDIT4* after cardiac arrest (Tissier et al., 2019). In our study, these genes were upregulated in monocyte cluster 4, suggesting that our scRNA-seq dataset may be a useful resource for provisional “deconvolution” of whole blood transcriptomic datasets of cardiac arrest. That is, the scRNA-seq dataset (e.g., via our browser-based visualization https://viz.stjude.cloud/chen-lab/visualization/single-cell-transcriptomics-reveal-a-hyperacute-cytokine-and-immune-checkpoint-axis-in-patients-with-poor-neurological-outcomes-after-cardiac-arrest~204, https://tinyurl.com/54t86fuu) could be used to generate hypotheses for the cellular source of genes measured in whole blood.

We acknowledge several limitations of our study. Our analysis of PBMC does not examine end-organs, which have not been immunophenotyped in clinical CA. Experimental IRI models have highlighted the infiltration of innate immune cells to end-organs, and suggest a hypothesis that the cell subpopulations in our study represent immune cells in transit to end-organs. In animal models of CA, monocytes expand in circulation and accumulate in the brain in areas of neuronal injury (Giuliano et al., 2021; Jiang et al., 2020; Uray et al., 2021; Xing and Lu, 2017; Zhang et al., 2018). In experimental single-organ IRI, activated monocytes and NK cells migrated to sites of injury early in myocardial IRI (Bajpai et al., 2019), stroke (Chu et al., 2015; Planas, 2018; Zhang et al., 2014), and renal IRI (Zhang et al., 2020, 2008). Thus, we defer to future studies to adjudicate whether the early innate immune endotypes represent cells in transit to end-organs and define the role of tissue-resident cells. A second limitation is the small number of CA patients examined by scRNA-seq, as viable PBMC suitable for single-cell studies were previously not part of most CA biorepositories.

However, scRNA-seq studies of other human diseases have yielded valuable findings despite similar sample sizes (Reyfman et al., 2019). Further, since our scRNA-seq dataset was intended to be hypothesis-generating, we confirmed findings in two independent, validation cohorts at the cellular, protein and global transcriptomic level. The limited sample size of scRNA-seq prevented association with patient characteristics, such as etiology of CA; however, it could be argued that the consistency of the results across cohorts without accounting for patient heterogeneity or treatment differences supports that our findings are robust. Neutrophils were not included in this study due to the technical challenges of their cryopreserving neutrophils for droplet-based sequencing or running single-cell studies on fresh samples from CA patients presenting at irregular intervals (which would exacerbate batch effects in scRNA-seq).

In conclusion, the transcriptomic analysis of clinical OHCA patients at single-cell resolution revealed that, as early as 6h post-arrest, innate immune networks distinguished the neurological outcome at hospital discharge, even though this clinical outcome may not be apparent for days after hospital admission. Dysregulated innate immune cell subpopulations expressing immune checkpoint receptors associated with poor neurological outcomes. In a negative feedback loop, the pro- and anti-inflammatory cytokine environment induced expression of Nectin-2 on monocytes, which in turn limited production of proinflammatory cytokines by Tim-3^+^TIGIT^+^ NK cells. These acutely emerging and transient immunophenotypes highlight the coexistence of inflammation with endogenous, immunoregulatory mechanisms. Early augmentation of anti-inflammatory pathways may be a potential therapeutic time window and target for mitigating systemic inflammation after CA. Together, these data further our understanding of the immune landscape in CA and define approaches to targeted immunotherapy.

## Acknowledgments

Adam T. Chicoine (BWH Center for Cellular Profiling) provided flow cytometry expertise, and Sam J. Saliba provided multiplex plasma analysis expertise. This study was supported by the Japan Heart Foundation/Bayer Yakuhin Research Grant Abroad (T.T.), Zoll Foundation (T.T.), American Lebanese Syrian Associated Charities (XC), American Heart Association (AHA) 20TPA35500016 (E.Y.K.), AHA 15FTF25080205, Brigham and Women’s Hospital Department of Medicine Evergreen Innovation Fund (E.Y.K.). A.J.W. was supported by Zoll Foundation. We are grateful to other members of the Registry of Critical Illness who assisted with initial patient recruitment and phenotyping at BWH (Laura Fredenburgh, M.D., Sam Ash, M.D., Paul Dieffenbach, M.D.).

## Author contributions

This study was conceived by E.Y.K. and designed by E.Y.K., T.T., C.C., X.C.. The overall study was supervised by E.Y.K., X.C.. The clinical enrollment and biospecimen banking were performed by T.T., M.A.P.-V., A.J.W., E.A.B., K.M.B., and supervised by P.C.H., R.R.S., R.M.B., E.A.B., D.A.M., E.Y.K. Clinical chart review was done by T.T, J.V., M.A.P.-V., and supervised by R.M.B. and E.Y.K. The multiplex assays were designed, performed by T.T., S.J.S., and supervised by Y.T.. In vitro experiments were performed by T.T. and supervised by E.Y.K.. Bioinformatics analyses were performed by C.C., W.C. and supervised by X.C.. The manuscript was written by T.T. and C.C., and critical revision was made by L.T.M., W.M.O., P.R.L., D.A.M., F.I., X.C., E.Y.K.. Project administration by E.Y.K.. Funding acquisition by E.Y.K. and T.T. All authors have approved the submitted version.

## METHODS

### Study design and patient cohorts

The clinical discovery cohort of this study was a prospective, single-center observational investigation conducted at Brigham and Women’s Hospital (BWH). Validation studies were performed at both a single-center (BWH: flow cytometry, bulk RNA-seq) and multi-center (BWH and Beth Israel Deaconess Medical Center [BIDMC]: observational study, BWH). This study was approved by the IRB of each institution (Protocol ID: 2008P000495, 2014P002558). We prospectively enrolled adult (age ≥18 years) patients who were successfully resuscitated after out-of-hospital cardiac arrest (OHCA) between January 2014 and March 2020. Written consent was obtained from the patient or a legal surrogate if the patient did not achieve full recovery of consciousness (Glasgow Coma Scale [GCS] of 15) and competence for medical decision-making. All patients were treated in accordance with the latest post-arrest guidelines (Callaway et al., 2015). Written consent was obtained from non-CA control patients and healthy subjects. Post-OHCA patients were categorized as having a good neurological outcome (“good outcome”) or poor neurological outcome (“poor outcome”) by Cerebral Performance Category (CPC) at hospital discharge. Good and poor neurological outcome was defined as CPC score 1 to 2 (*i.e.*, independent activities of daily life [ADL]), and CPC 3 to 5 (*i.e.*, ranging from conscious but dependent for ADL [3] to coma [4] or brain death [5]), respecitvely (Becker et al., 2011). The following variables were recorded for post-CA patients: age, sex, location of CA, witness status, implementation of bystander cardiopulmonary resuscitation (CPR), initial heart rhythm, presumed duration of CA, etiology of arrest (cardiac or non-cardiac), comorbidities, and CPC at hospital discharge.

### Blood sample processing

Peripheral blood was collected in a blood collection tube containing ethylenediaminetetraacetic acid [EDTA] (Becton, Dickinson and Company). Plasma was isolated by centrifugation at 2,000 × *g* for 20 min and stored at -80°C. Peripheral blood mononuclear cells (PBMC) were isolated using the density gradient centrifugation method (GE Healthcare, STEMCELL Technology). Briefly, gradients were centrifuged at 1,200 × *g* for 20 min at room temperature. The PBMC interface was centrifuged and washed and cryopreserved until further analyses.

### Single-cell RNA-sequencing and Preprocessing

PBMC from 11 post-OHCA patients and 3 healthy subjects were used. Among the 11 OHCA patients, 5 (2 with good outcomes and 3 with poor outcomes) have been profiled at both time points (6h and 48h), 2 at 6-hour only (1 good and 1 poor outcome) and 4 at 48-hours only (1 good and 3 poor outcomes). The single-cell suspensions of scRNA-seq samples were converted to barcoded scRNA-seq libraries using the Chromium Single Cell 3′ Library, Gel Bead and Multiplex Kit, and Chip Kit (10X Genomics). The Chromium Single Cell 3′ v2 Reagent (10X Genomics, 120237) kit was used to prepare single-cell RNA libraries according to the manufacturer’s instructions. Cell Ranger Single-Cell Software Suite (version 2.1.1, 10X Genomics) was used to quality control the single-cell expression data. The filtered gene-barcode matrix output from Cell Ranger, was analyzed in further analyses using Clustergrammer275. UMIs mapped to genes encoding ribosomal/mitochondrial proteins were removed, and cells with more than 60% of UMIs mapped to ribosomal/mitochondrial proteins were filtered. Cells with low (≤1,024) or high (≥16,384) UMI counts were further filtered. A total of 104,609 cells were captured with an average of 4,214 mRNA molecules (UMIs, median: 3,300, range: 1,025-16,383). Each gene’s expression is normalized to 10,000 UMIs per cell and log-transformed by adding 1 to the expression matrix.

Transcriptomes from each outcome cohort and time point (18,685 cells in healthy, 20,031 cells in 6h-good outcome, 16,039 cells in 6h-poor outcome, 16,707 cells in 48h-good outcome, and 24,717 cells in 48h-poor outcome) were subjected to quality control filtering and merged into a single data set of 96,179 individual cells.

### scRNA-seq analysis

<u>Clustering</u> The whole dataset’s subpopulation structure was inferred using Latent Cellular State Analysis (Cheng et al., 2019). The optimal number of clusters was manually selected from top models determined by the silhouette measure for solutions with different clusters (from 2 to 30). <u>Data visualization.</u> High dimensional scRNA-seq data were visualized on two-dimensional maps through t-distributed stochastic neighbor embedding (t-SNE) using R language with default settings. Patterns of gene expression for cell types and gene groups were visualized using R package *ComplexHeatmap* (Gu et al., 2016). General visualizations were performed using packages *ggplot2* and *ggpubr*. <u>Differential gene expression analysis.</u> Differential expressions between different conditions or cell types were analyzed by the negative binomial with independent dispersions (Chen et al., 2018) with batch effect inferred using SVA (Chen et al., 2020). <u>Pathway analysis.</u> Gene set enrichment analysis and pathway analysis were performed using R package *ClusterProfiler* (Yu et al., 2012). We used Molecular Signature Database for gene annotations, including Hallmark gene sets. GSEA results were visualized using R package *enrichplot*.

<u>Interactome analysis.</u> NicheNet was used to prioritize the ligand-receptor pairs predicted to be involved in interactions among cell types (Browaeys et al., 2020). We input the signatures of differentially expressed genes between patients with poor and good neurological outcomes in sender and receiver cells. We selected the predominant cell population of each outcome to represent the corresponding conditions to compare the different cell networks among neurological outcomes. Namely, for NK cells, we used cluster 2 of the patients with good outcomes at 6h-post CA as the reference condition and cluster 1 of the patients with poor outcomes as the condition of interest. Likewise, for monocytes, we used cluster 2 and cluster 4 of good and poor outcomes at 6h post-CA, respectively. The differentially expressed genes in scRNA-seq were identified using NBID and SVA. <u>M1/M2 signature score.</u> Monocytes/macrophages are often classified as classically activated proinflammatory M1 phenotype or alternatively activated M2 phenotype which is pro-resolution of inflammation and involved in the repair of damaged tissues. We used the literature to curate a list of genes that were previously described as M1 and M2 signature genes (**Supplemental Data File 3**) and calculated the M1/M2 score for clusters 2 and 4 by averaging the expression levels of these genes in each cell.

### Flow cytometry and cell sorting

PBMC from 12 healthy subjects and 28 post-CA patients (N = 12 and N = 16, for good and poor outcomes, respectively) were analyzed by flow cytometry (**Supplemental Table 2**). Among these subjects, Tim-3^+^ NK cells were sorted from post-CA patients with poor outcomes at 6h-CA (N = 8). Tim-3^-^ NK cells were sorted from healthy subjects and patients with good outcomes at 6h post-CA (N = 11, N = 5 respectively). Nectin-2^+^ monocytes were sorted from patients with poor outcomes at 6h post-CA (N = 10). Nectin-2^-^ monocytes were sorted from both healthy subjects and patients with good outcomes at 6h post-CA (N = 11, N = 10, respectively). PBMC were incubated with Fc receptor-blocking antibody (eBioscience) and viability dye (eBioscience) in 1x PBS for 15 min at 4 °C. Cells were then incubated with human-specific antibodies in a staining buffer (PBS with 0.5% FBS and 0.5mM EDTA) for 30 min at 4°C. Cells were sorted with 4-laser BD FACSAria™ Fusion cell sorter (BD Life Sciences). Data was acquired using FACSDiva software^TM^ and analyzed by FlowJo 10.7.1 software (all from Becton, Dickinson and Company). Cell subsets were defined by FSC-A/SSC-A, singlets by FSC-A/FSC-H, dead cells were gated out with viability dye (eBioscience), and as follows: Tim3^+^ NK cells (CD3^-^CD15^-^CD14^-^CD56^+^Tim3^+^), Tim3^-^ NK cells (CD3^-^CD15^-^CD14^-^CD56^+^Tim3^-^), Nectin2^+^ monocytes (CD3^-^CD15^-^CD14^+^Nectin2^+^), Nectin2^-^ monocytes (CD3^-^CD15^-^CD14^+^Nectin2^-^). The following anti-human antibodies were purchased from Biolegend: anti-CD3 (HIT3A), anti-CD14 (M5E2), anti-CD15 (H198), anti-CD16 (3G8), anti-CD56 (5.1H11), anti-CD366 (Tim-3) (F38-2E2), and anti-Nectin-2 (TX31).

### Bulk RNA sequencing and analysis

Sorted PBMC were lysed in TCL buffer (Qiagen) + 1% beta-mercaptoethanol. Subsequently, RNA extraction and library preparation and sequencing were done as previously described (Kim et al., 2020). Bulk RNA data were processed using workflow for the analysis of RNA-Seq differential expression (WARDEN) (McLeod et al., 2021). FastQ files generated by RNA-Seq are mapped to the reference genome (GRCh37) using the RNA-seq aligner STAR. The average mapping rate is 84.68%, with an average of 11.20 million mapped reads per sample. Read counts per gene were analyzed using R software for differential expression analysis. Low count genes were removed from analysis using a CPM cutoff corresponding to a count of 10 reads.

Normalization factors were generated using the TMM method. Counts were normalized using voom. Voom normalized counts were analyzed using the lmFit and eBayes functions of the limma software package. We selected the top 500 genes with the smallest adjusted P values from the DE analysis of NK cells and monocytes in either scRNA-seq and bulk RNA-seq to avoid noise caused by weak signals and the fold changes from each method were compared using Spearman correlation.

### Plasma validation

Plasma samples were prospectively collected from 51 OHCA patients who were admitted to BWH and BIDMC after successful resuscitation (N = 18, N = 30, good or poor outcomes, respectively) and non-CA control patients and healthy volunteers in the BWH plasma biobank (N = 15) (**Supplemental Table 3**). Plasma levels of cytokines (IFN-γ, TNFα, IL-10, IL-18, and CCL2) were measured using the Luminex MAGPIX system (Luminex) and commercially available kit (MILLIPLEX) as per the manufacturer’s instruction.

### Human in vitro model

<u>Monocyte stimulation.</u> CD14^+^ monocytes were isolated from PBMC from healthy subjects by positive selection withanti-CD14 magnetic labeling beads and an isolation column (Miltenyi Biotec). Monocytes were resuspended in RPMI 1640 medium (Gibco) supplemented with 2% FBS, and incubated at 5 × 10^4^ cells per well in a flat-bottom 96-well plate (Corning) for 6h with stimulants.

The stimulants screened included recombinant human TNFα 50ng/mL (210-TA/CF), IFN-γ 100ng/mL (285-IF-100/CF), IL-18 100ng/mL (9124-IL-010/CF), IL-10 100ng/mL (217-IL-005/CF), IL-27 50ng/mL (2526-IL-010/CF), TGF-β1 20ng/mL (7754-BH-005/CF), LIGHT 100ng/mL (664-LI-025/CF), molidustat 20nM (Cayman), or vehicle (PBS + 0.5% DMSO). All recombinant human cytokines were purchased from R&D Systems. After 6h of incubation, gene expression levels were analyzed with RT-QPCR.

<u>Incubation of PBMC from healthy volunteer with anti-Nectin-2 antibody.</u> Freshly isolated PBMC from healthy volunteers were incubated in the media (RPMI1640 + 2% FBS) with IFN-γ 100ng/mL and IL-10 100ng/mL in the presence or absence of human anti-Nectin-2 monoclonal antibody 20 μg/mL (R&D Systems) for 6h. After stimulation, phorbol myristate acetate (PMA), ionomycin (eBioscience), and brefeldin A (eBioscience) were added and further incubated for 4h.

### Incubation of PBMC from post-CA patients with anti-Nectin-2 antibody or recombinant human

<u>Nectin-2.</u> Frozen PBMC of post-CA patients were thawed and incubated in the media (RPMI1640 +2% FBS) with human anti-Nectin-2 monoclonal antibody or isotype control (20 μg/mL, both R&D Systems) for 6h. After the incubation, brefeldin A was added and incubated for 4h. Frozen PBMC of post-CA patients were thawed and incubated in media (RPMI1640 + 2% FBS) with recombinant human Nectin-2 (Biolegend) or isotype control (ENZO) at a concentration of 100 ng/mL for 6h.

After the incubation, PMA, ionomycin, and brefeldin A was added and incubated for 4h.

<u>Flow cytometry analysis.</u> The abundance of Nectin-2^+^ monocytes and NK cell intracellular cytokine production of IFN-γ (4S.B3, Biolegend) were analyzed by flow cytometry (BD LSRFortessa™ Cell Analyzer, BD Biosciences) as described previously.

### Quantitative PCR

RNA was extracted using RNeasy Mini Kits (Qiagen). cDNA was prepared using Quantitect RT-PCR (Qiagen) and RT Q-PCR was performed using the TaqMan Gene Expression Assay (ThermoFisher Scientific) on an AriaMx (Agilent). The same quantity of cDNA was put into each reaction, and relative expression was normalized to the housekeeping gene UBC. Relative expression was calculated by normalizing to the unstimulated control condition of each subject. Optimized primer/probes sets were purchased from ThermoFisher Scientific with the following assay IDs: NECTIN2 (Hs01071562_m1), TNF (Hs00174128_m1), IL12A (Hs01073447_m1), IL18 (Hs01038788_m1), IL10 (Hs00961622_m1), TGFB1 (Hs00998133_m1), and UBC (Hs00824723_m1).

### Statistical analysis

For continuous variables, normally distributed data are represented as mean ± SD, and skewed data are shown as median with interquartile range. Cell type or cluster abundance is shown as median with range. For RNA sequencing experiments, see the detailed methods above under “Single-cell RNA-sequencing” and “Bulk RNA sequencing”. Mixed effect logistic regression was used for testing the difference of cell-type frequencies among different patient groups. The data process and test were done by R package lme4 (Yang et al., 2018). For flow cytometry analysis, ANOVA with Tukey’s multiple comparisons or Kruskal-Wallis test with Dunn’s multiple comparisons was used as appropriate for three-group comparisons. For a two-group comparison, unpaired Student t-test or Mann-Whitney test was used as appropriate. Benjamini-Hochberg method was used for the adjustment for multiple comparisons in the multiplex assay. A value of P <0.05 was considered statistically significant. All analyses other than RNA sequencing were done using GraphPad Prism 9.0.0 (GraphPad Software Inc., La Jolla, CA).

## List of Supplementary Figures

Figure S1. Graphical summary.

Figure S2. Cell type-defining genes and frequencies of major cell lineages.

Figure S3. Monocyte analysis after CA: proportions of classical and non-classical monocyte subsets do not distinguish CA patients with good or poor neurological outcome; genes and gene sets enriched in monocyte clusters post-CA in scRNA-seq.

Figure S4. Proportions of CD56^dim/bright^CD16^+/-^ NK cells do not distinguish CA patients with good or poor neurological outcome.

Figure S5. Validation cohort confirms the Nectin-2^+^ monocyte and Tim-3^+^ NK cell subpopulations identified by scRNA-seq in discovery cohort.

Figure S6. IFNg and IL-10 do not cooperate in stimulation of cytokine production by monocytes.

## List of Supplementary Tables

**Table S1.**
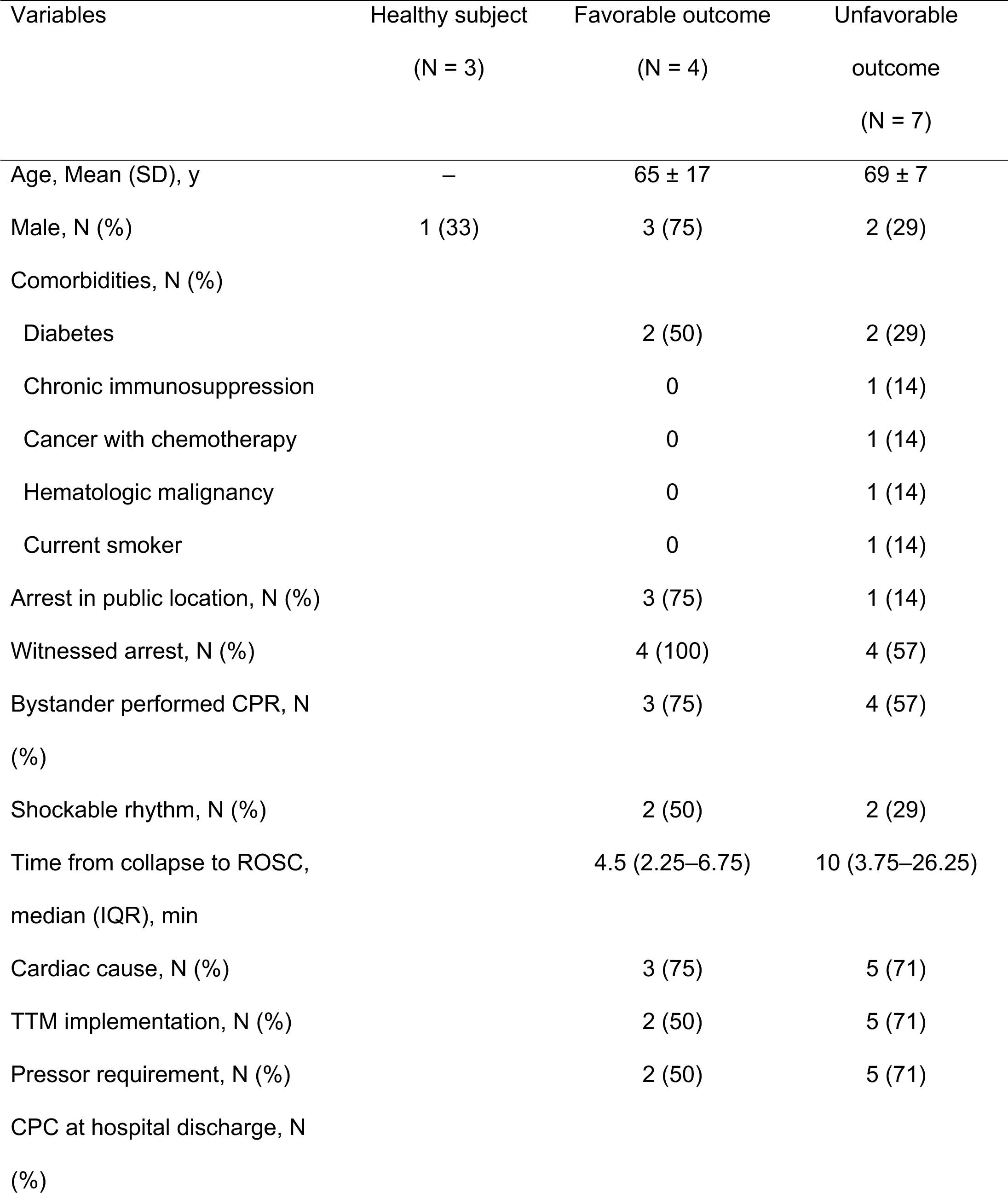
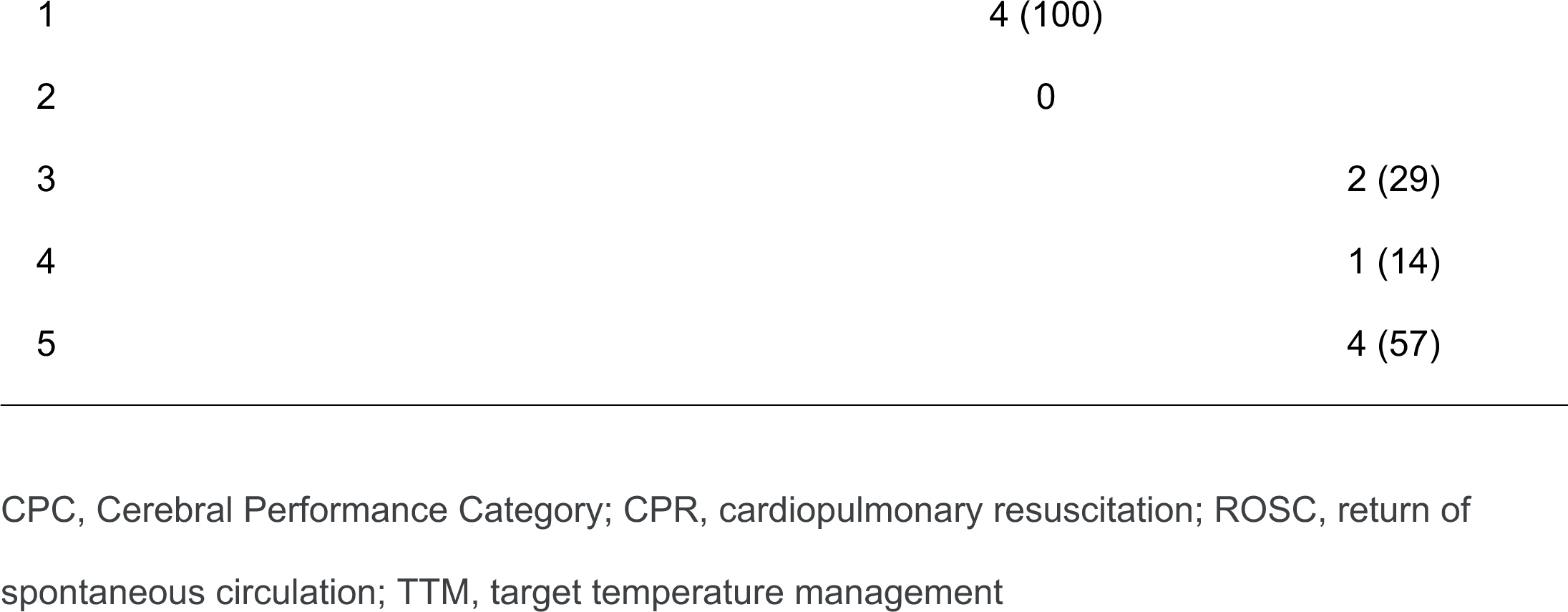
Characteristics of participants in single-cell RNA-sequencing analysis.

| Variables | Healthy subject<br>(N = 3) | Favorable outcome<br>(N = 4) | Unfavorable<br>outcome<br>(N = 7) |
| --- | --- | --- | --- |
| Age, Mean (SD), y | – | 65 ± 17 | 69 ± 7 |
| Male, N (%) | 1 (33) | 3 (75) | 2 (29) |
| Comorbidities, N (%) |  |  |  |
| Diabetes |  | 2 (50) | 2 (29) |
| Chronic immunosuppression |  | 0 | 1 (14) |
| Cancer with chemotherapy |  | 0 | 1 (14) |
| Hematologic malignancy |  | 0 | 1 (14) |
| Current smoker |  | 0 | 1 (14) |
| Arrest in public location, N (%) |  | 3 (75) | 1 (14) |
| Witnessed arrest, N (%) |  | 4 (100) | 4 (57) |
| Bystander performed CPR, N<br>(%) |  | 3 (75) | 4 (57) |
| Shockable rhythm, N (%) |  | 2 (50) | 2 (29) |
| Time from collapse to ROSC,<br>median (IQR), min |  | 4.5 (2.25–6.75) | 10 (3.75–26.25) |
| Cardiac cause, N (%) |  | 3 (75) | 5 (71) |
| TTM implementation, N (%) |  | 2 (50) | 5 (71) |
| Pressor requirement, N (%) |  | 2 (50) | 5 (71) |
| CPC at hospital discharge, N<br>(%) |  |  |  |

Table S2. List of M1/M2 monocyte signature genes. (Excel file)

Table S3. List of scRNA-seq monocyte cluster-defining genes. (Excel file)

Table S4. List of DE genes of scRNA-seq monocyte cluster 4 vs. cluster 2. (Excel file)

Table S5. List of scRNA-seq NK cell cluster-defining genes. (Excel file)

Table S6. List of DE genes of scRNA-seq NK cell cluster 1 vs. cluster 2. (Excel file)

**Table S7.**
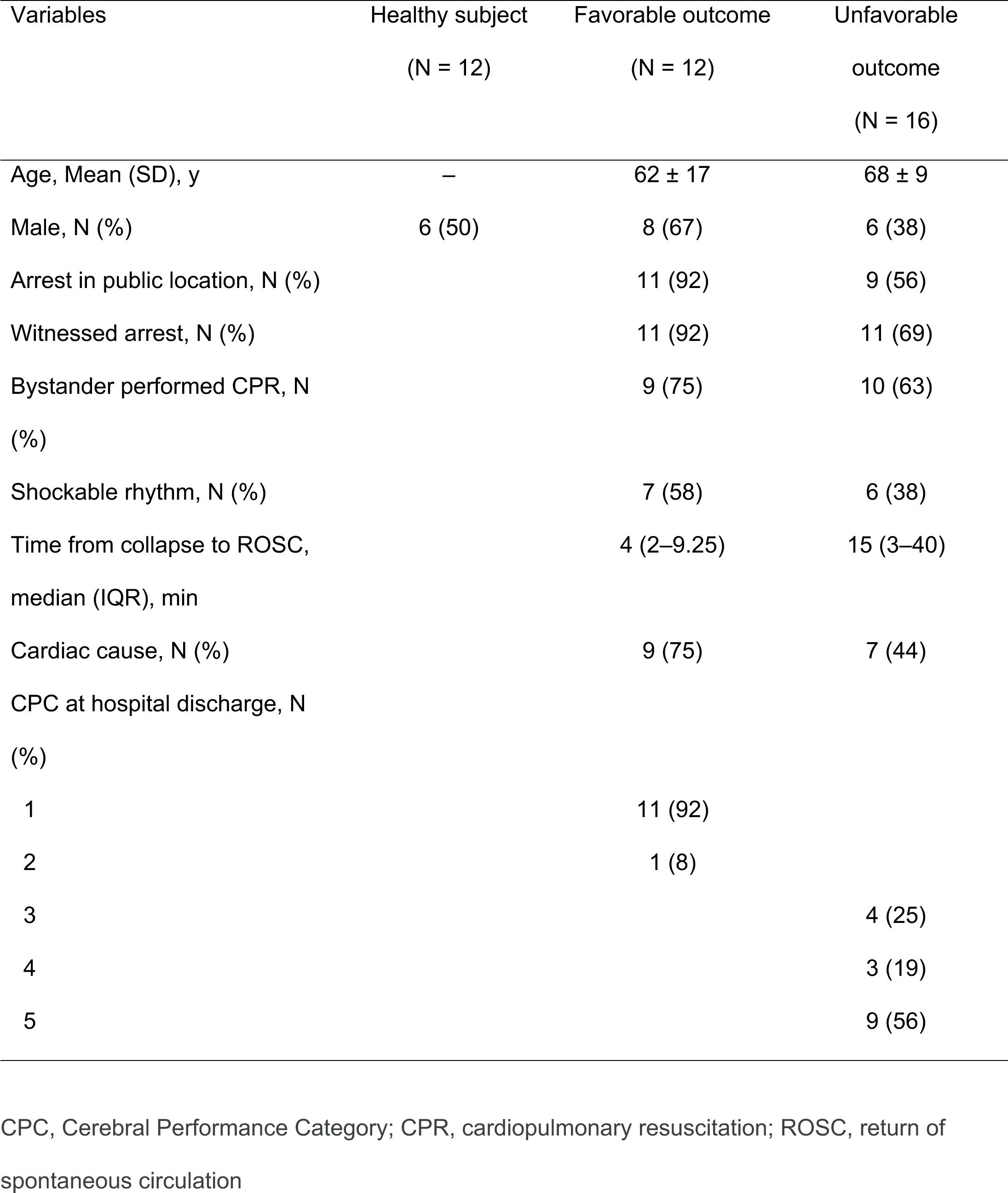
Characteristics of participants in flow cytometry analysis.

| Variables | Healthy subject<br>(N = 12) | Favorable outcome<br>(N = 12) | Unfavorable<br>outcome<br>(N = 16) |
| --- | --- | --- | --- |
| Age, Mean (SD), y | – | 62 ± 17 | 68 ± 9 |
| Male, N (%) | 6 (50) | 8 (67) | 6 (38) |
| Arrest in public location, N (%) |  | 11 (92) | 9 (56) |
| Witnessed arrest, N (%) |  | 11 (92) | 11 (69) |
| Bystander performed CPR, N<br>(%) |  | 9 (75) | 10 (63) |
| Shockable rhythm, N (%) |  | 7 (58) | 6 (38) |
| Time from collapse to ROSC,<br>median (IQR), min |  | 4 (2–9.25) | 15 (3–40) |
| Cardiac cause, N (%) |  | 9 (75) | 7 (44) |
| CPC at hospital discharge, N<br>(%) |  |  |  |
| 1 |  | 11 (92) |  |
| 2 |  | 1 (8) |  |
| 3 |  |  | 4 (25) |
| 4 |  |  | 3 (19) |
| 5 |  |  | 9 (56) |

CPC, Cerebral Performance Category; CPR, cardiopulmonary resuscitation; ROSC, return of spontaneous circulation

Table S8. List of DE genes of bulk RNA-seq Nectin-2^+^ vs. Nectin-2^-^ monocytes. (Excel file)

Table S9. List of DE genes of bulk RNA-seq Tim-3^+^ vs Tim-3^-^ NK cells. (Excel file)

**Table S10.** Characteristics of participants in plasma cytokine/chemokine assay.

| Variables | Healthy subject<br>(N = 15) | Favorable outcome<br>(N = 18) | Unfavorable<br>outcome<br>(N = 30) |
| --- | --- | --- | --- |
| Age, Mean (SD), y | 68 ± 12 | 66 ± 17 | 69 ± 12 |
| Male, N (%) | 8 (53) | 9 (50) | 15 (50) |
| Arrest in public location, N (%) |  | 16 (89) | 12 (41) |
| Witnessed arrest, N (%) |  | 18 (100) | 19 (65) |
| Bystander performed CPR, N<br>(%) |  | 15 (83) | 19 (65) |
| Shockable rhythm, N (%) |  | 12 (67) | 9 (30) |
| Time from collapse to ROSC,<br>median (IQR), min |  | 3 (2–9) | 15 (9–37.5) |
| Cardiac cause, N (%) |  | 13 (72) | 7 (24) |
| CPC at hospital discharge, N<br>(%) |  |  |  |
| 1 |  | 15 (83) |  |
| 2 |  | 3 (17) |  |
| 3 |  |  | 2 (7) |
| 4 |  |  | 2 (7) |
| 5 |  |  | 26 (87) |

CPC, Cerebral Performance Category; CPR, cardiopulmonary resuscitation; ROSC, return of spontaneous circulation

