## Supplementary material for "Single-cell transcriptomics reveal hyperacute cytokine and immune checkpoint axis in patients with poor neurological outcomes after cardiac arrest": Supplemantal Information

Tamura T, Cheng C, *et. al*.

List of Supplementary Figures

Figure S1. Graphical summary.
Figure S2. Cell type-defining genes and frequencies of major cell lineages.
Figure S3. Monocyte analysis after CA: proportions of classical and non-classical monocyte subsets do not distinguish CA patients with good or poor neurological outcome; genes and gene sets enriched in monocyte clusters post-CA in scRNA-seq.
Figure S4. Proportions of CD56$^{dim/bright}$CD16$^{+/-}$ NK cells do not distinguish CA patients with good or poor neurological outcome.
Figure S5. Validation cohort confirms the Nectin-2+ monocyte and Tim-3+ NK cell subpopulations identified by scRNA-seq in discovery cohort.
Figure S6. IFNg and IL-10 do not cooperate in stimulation of cytokine production by monocytes.

List of Supplementary Tables

Table S1. Characteristics of participants in single-cell RNA-sequencing analysis.
Table S2. List of M1/M2 monocyte signature genes. (Excel file)
Table S3. List of scRNA-seq monocyte cluster-defining genes. (Excel file)
Table S4. List of DE genes of scRNA-seq monocyte cluster 4 vs. cluster 2. (Excel file)
Table S5. List of scRNA-seq NK cell cluster-defining genes. (Excel file)
Table S6. List of DE genes of scRNA-seq NK cell cluster 1 vs. cluster 2. (Excel file)
Table S7. Characteristics of participants in flow cytometry analysis.
Table S8. List of DE genes of bulk RNA-seq Nectin-2$^+$ vs. Nectin-2$^-$ monocytes. (Excel file)
Table S9. List of DE genes of bulk RNA-seq Tim-3$^+$ vs Tim-3$^-$ NK cells. (Excel file)
Table S10. Characteristics of participants in plasma cytokine/chemokine assay.

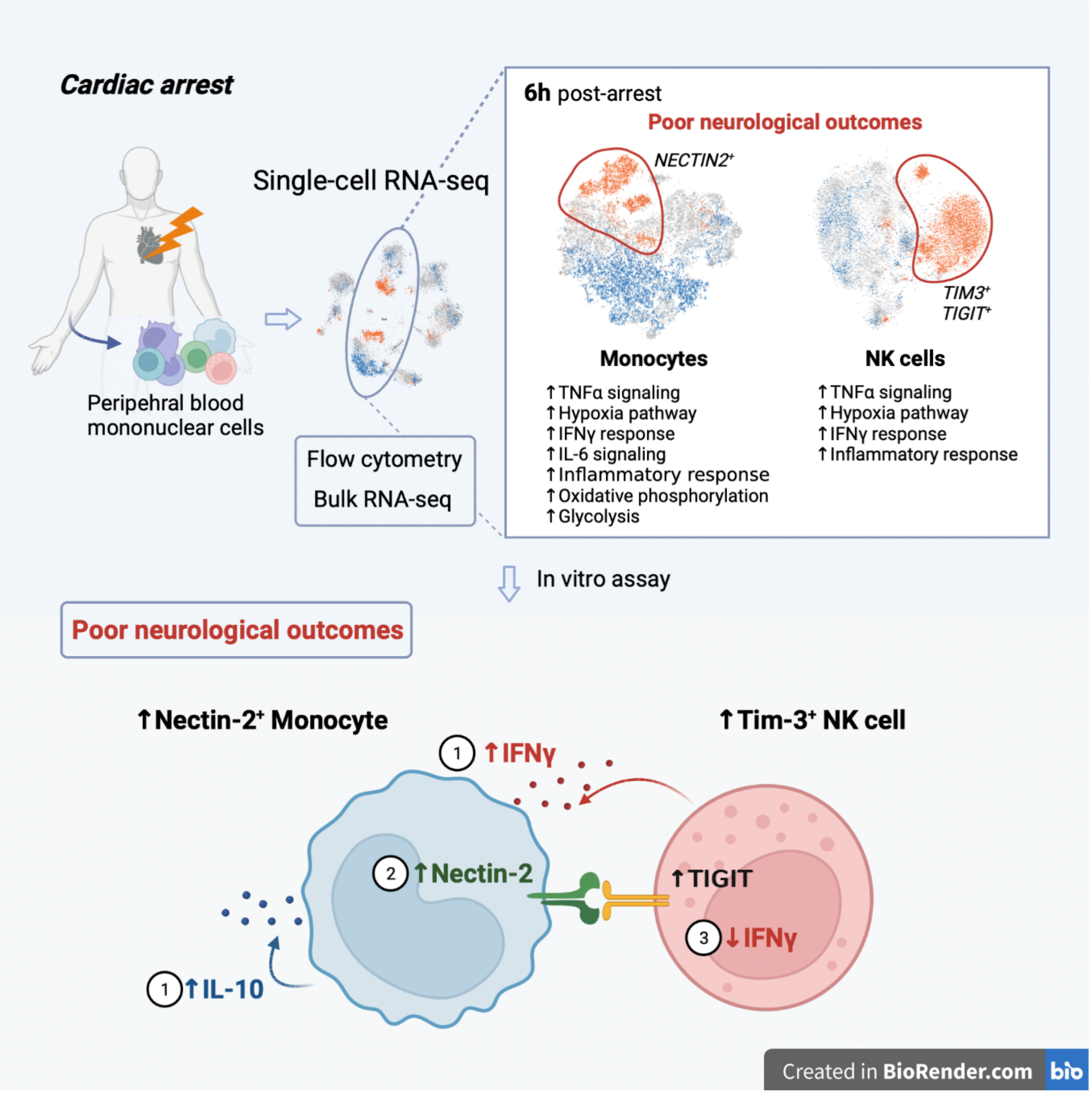


**Figure S1. Graphical summary.** Single-cell RNA-sequencing revealed unique monocyte and NK cell states associated with poor neurological outcomes after cardiac arrest. Ex vivo studies on PBMC from post-arrest patients established a scheme in which the mixed pro- and anti-inflammatory cytokine milieu induces expression of Nectin-2$^+$ on monocytes. Nectin-2 then drives a negative feedback loop to limit inflammation by reducing IFNγ production by NK cells.

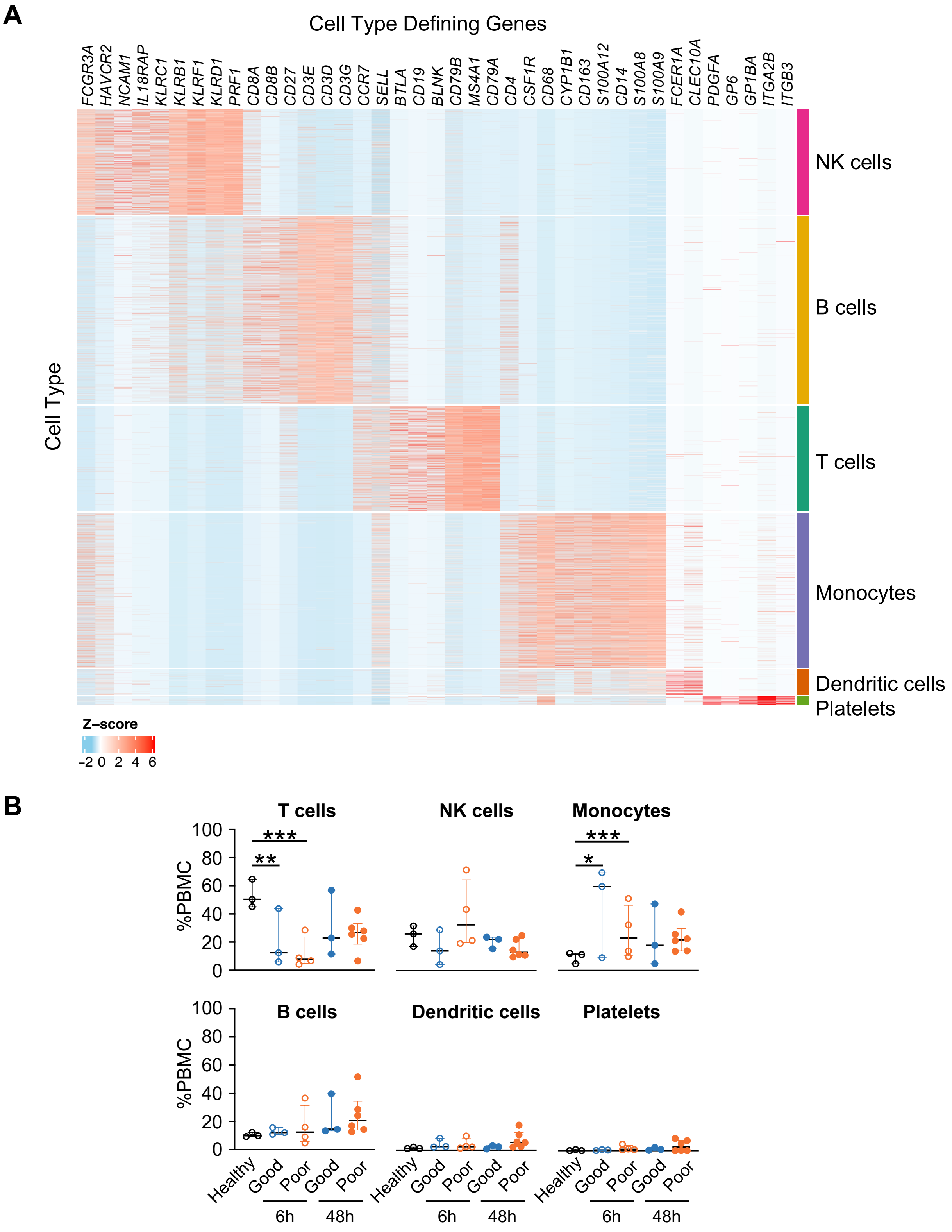


**Figure S2. Cell type-defining genes and frequencies of major cell lineages. A)** Heatmap of cell type-defining genes for major cell lineages is shown for the scRNA-seq dataset of PBMC from healthy subjects or patients post-CA (Figure 1). CD14 or CD163 was not highlighted in platelets–monocyte aggregates since row normalization of gene expression was performed. **B)**

Frequency of major cell lineages are shown per patient. NK, natural killer. B, mixed effect logistic regression, *P <0.05, **P < 0.01, ***P < 0.001.

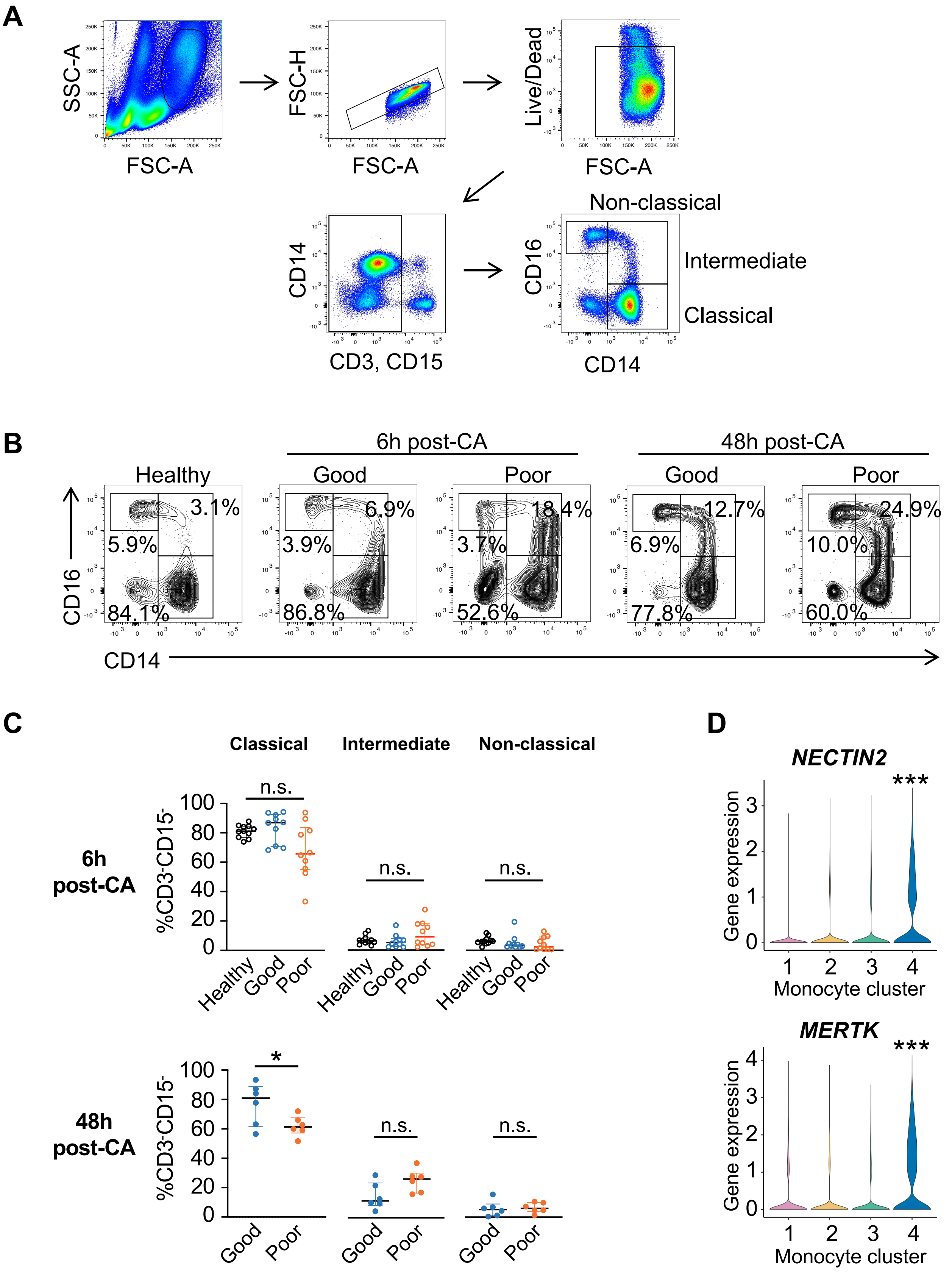
A
SSC-A
FSC-A
FSC-H
FSC-A
Live/Dead
FSC-A
CD14
CD3, CD15
CD16
CD14
Non-classical
Intermediate
Classical
B
6h post-CA
48h post-CA
Healthy
Good
Poor
Good
Poor
3.1%
5.9%
84.1%
6.9%
3.9%
86.8%
18.4%
3.7%
52.6%
12.7%
6.9%
77.8%
24.9%
10.0%
60.0%
CD16
CD14
C
Classical
Intermediate
Non-classical
6h post-CA
48h post-CA
%CD3-CD15-
n.s.
*
Healthy
Good
Poor
D
NECTIN2
MERTK
***
Gene expression
Monocyte cluster

**E**

TNFα signaling (Mono4 vs other Mono)

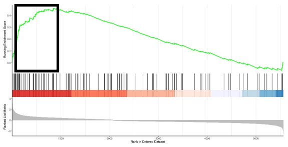

*AREG, GPR183, PHLDA1, CCL4, CCL2, CXCL3, MARCKS, ETS2, SDC4, CXCL2, LAMB3, DUSP4, KYNU, MAP3K8, SPHK1, NAMPT, LITAF, JUN, TNFAIP3, CD69, SOCS3, ATP2B1, MXD1, IL7R, FOSL1, GADD45B, SERPINE1, TLR2, PLPP3, ABCA1, ICOSLG, KLF9, PER1, SAT1, MYC, SOD2, NFKBIA, PFKFB3, MAFF, CCRL2, NR4A2, BCL3, TRIP10, RELB*

Inflammatory response (Mono4 vs other Mono)

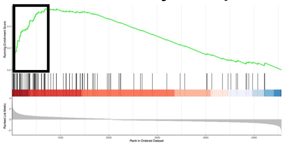

*GPR183, IL1R1, CXCL8, EREG, ADM, CCL2, TIMP1, CCL7, IL18R1, MSR1, HIF1A, SPHK1, MARCO, NAMPT, CMKLR1, MMP14, AHR, ITGB8, CD69, RGS, PTGER2, GNA15, CD14, ATP2B1, MXD1, IL7R, AQP9, PTAFR, SERPINE1, C5AR1, BEST1, TLR2, CD82, ABCA1, ICOSLG, IL10, C3AR1, MYC, RASGRP1, FZD5, NFKBIA, SCN1B, SLAMF1*

**Figure S3. Monocyte analysis after CA: proportions of classical and non-classical monocyte subsets do not distinguish CA patients with good or poor neurological outcome; genes and gene sets enriched in monocyte clusters post-CA in scRNA-seq.**

Flow cytometry (A-C) or scRNA-seq and monocyte fine clustering analysis (D-E) was performed on PBMC from healthy subjects or patients post-CA. **A)** Gating strategy for monocytes. **B)** Representative flow cytometry plots of classical ($CD14^{+}CD16^{-}$), intermediate ($CD14^{+}CD16^{+}$), and non-classical ($CD14^{-}CD16^{+}$) monocytes. **C)** Quantification of monocytes. **D)** Violin plot of cluster-defining genes for monocyte cluster 4 in Figure 2D. Gene expression levels of cluster 4 were significantly higher by pair-wise comparison with other clusters. **E)** Enrichment plots are shown for TNFa signaling and inflammatory response from GSEA analysis of monocyte cluster 4 in Figure 2F. Good and poor indicates neurological outcomes after CA. CA, cardiac arrest. C, D, Kruskal-Wallis test or Mann-Whitney U test; D, Kruskal-Wallis test with Dunn's multiple comparisons; *P < 0.05, ***P < 0.001.

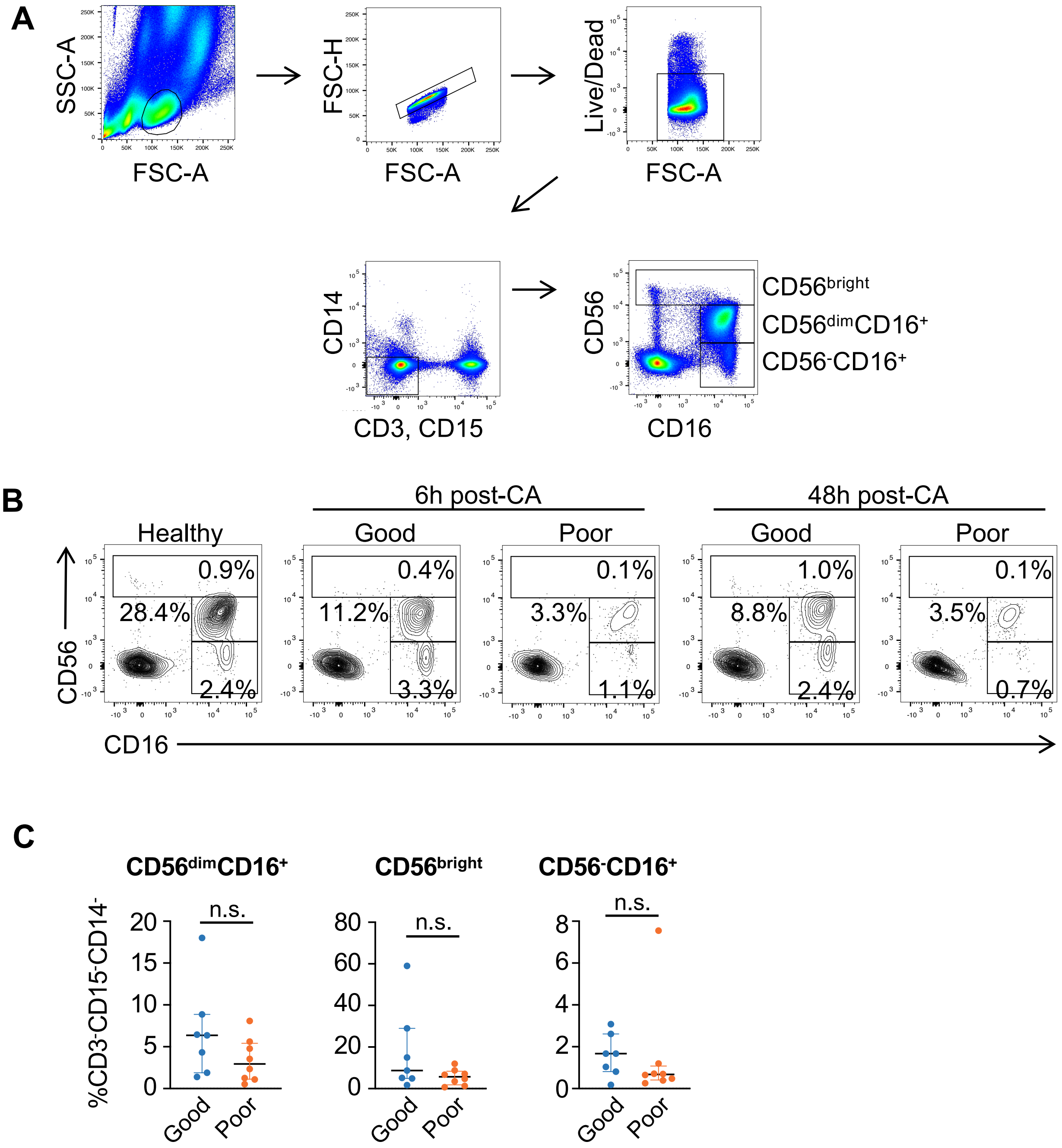


**Figure S4. Proportions of CD56$^{dim/bright}$CD16$^{+/-}$ NK cells do not distinguish CA patients with good or poor neurological outcome.** A-C) Flow cytometry was performed on PBMC from healthy subjects or patients post-CA. **A)** Gating strategy for NK cells. **B)** Representative flow cytometry plots of CD56$^{bright}$, CD56$^{dim}$CD16$^{+}$, and CD56$^{-}$CD16$^{+}$ NK cells. **C)** Quantification of NK cell subsets at 48h post-CA. C, Mann-Whitney U test. Good and poor indicates neurological outcomes after CA. CA, cardiac arrest; NK, natural killer.

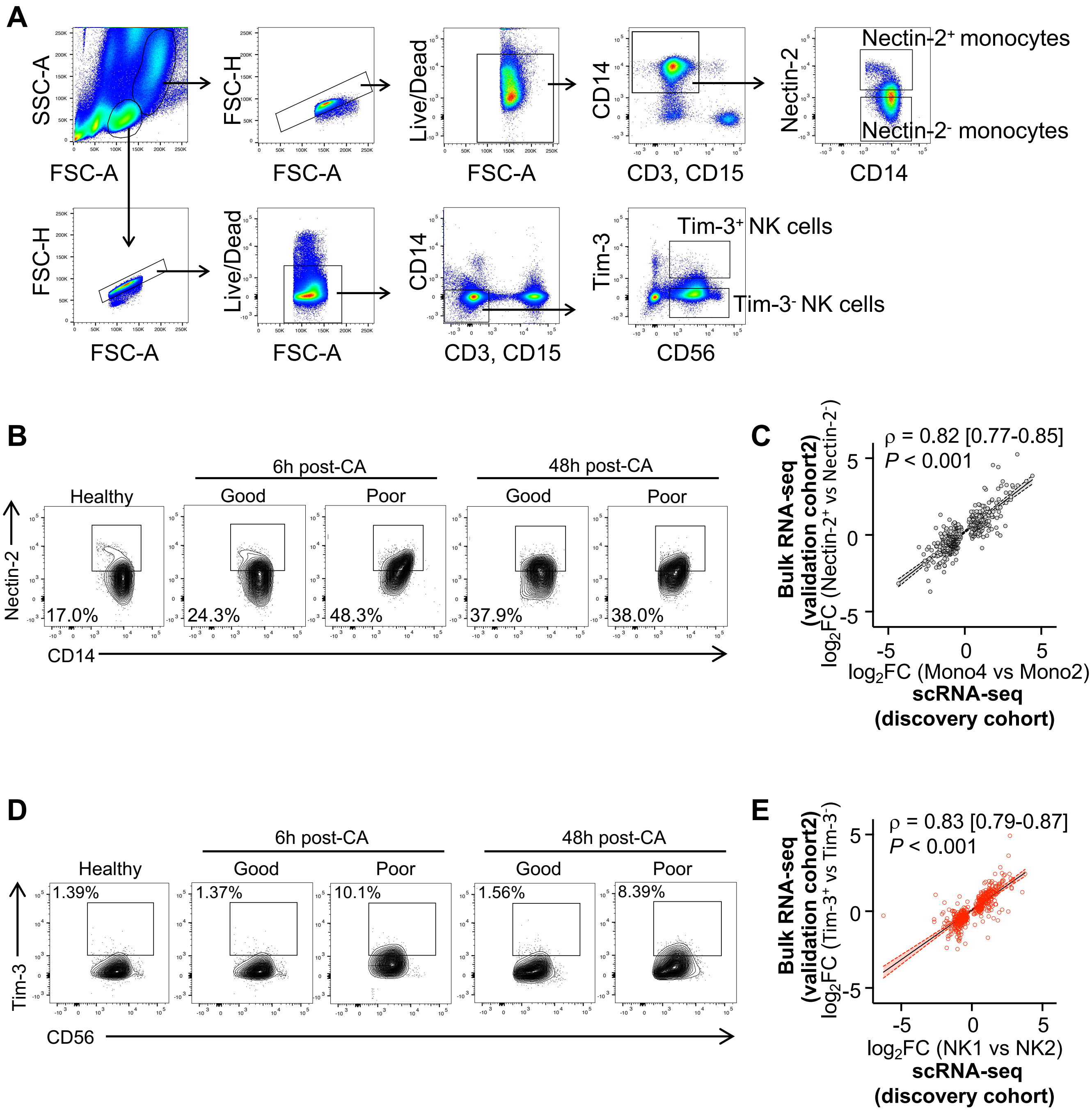


**Figure S5. Validation cohort confirms the Nectin-2+ monocyte and Tim-3+ NK cell subpopulations identified by scRNA-seq in discovery cohort. A)** Flow cytometric gating strategy for sorting of Nectin-$2^{+}$ and Nectin-$2^{-}$ monocytes, and Tim-$3^{+}$ Tim-$3^{-}$ NK cells. **B)** Flow cytometry analysis of Nectin-$2^{+}$ monocytes. Representative flow cytometry plots are shown. **C)** Nectin-$2^{+}$ and Nectin-2- $CD14^{+}$ monocytes were sorted by flow cytometry and assessed by bulk RNA-seq. DE of genes for Nectin-2+ monocyte cluster 4 compared with Nectin-2- monocyte cluster 2 were calculated for the scRNA-seq dataset. Correlation analysis of DE (FC) for monocyte scRNA-seq fine clustering compared to DE for bulk RNA-seq of sorted monocytes is shown. **D)** Flow cytometry analysis of Tim-$3^{+}$ NK cells.

Representative flow cytometry plots are shown. **E)** DE of genes for Tim-3$^{+}$ NK cell cluster 1 compared to Tim-3$^{-}$ NK cluster 2 were calculated for the scRNA-seq dataset. Correlation analysis of DE (FC) for NK cell scRNA-seq fine clustering compared to DE for bulk RNA-seq of sorted NK cells is shown. C, E, Spearman rank correlation analysis.

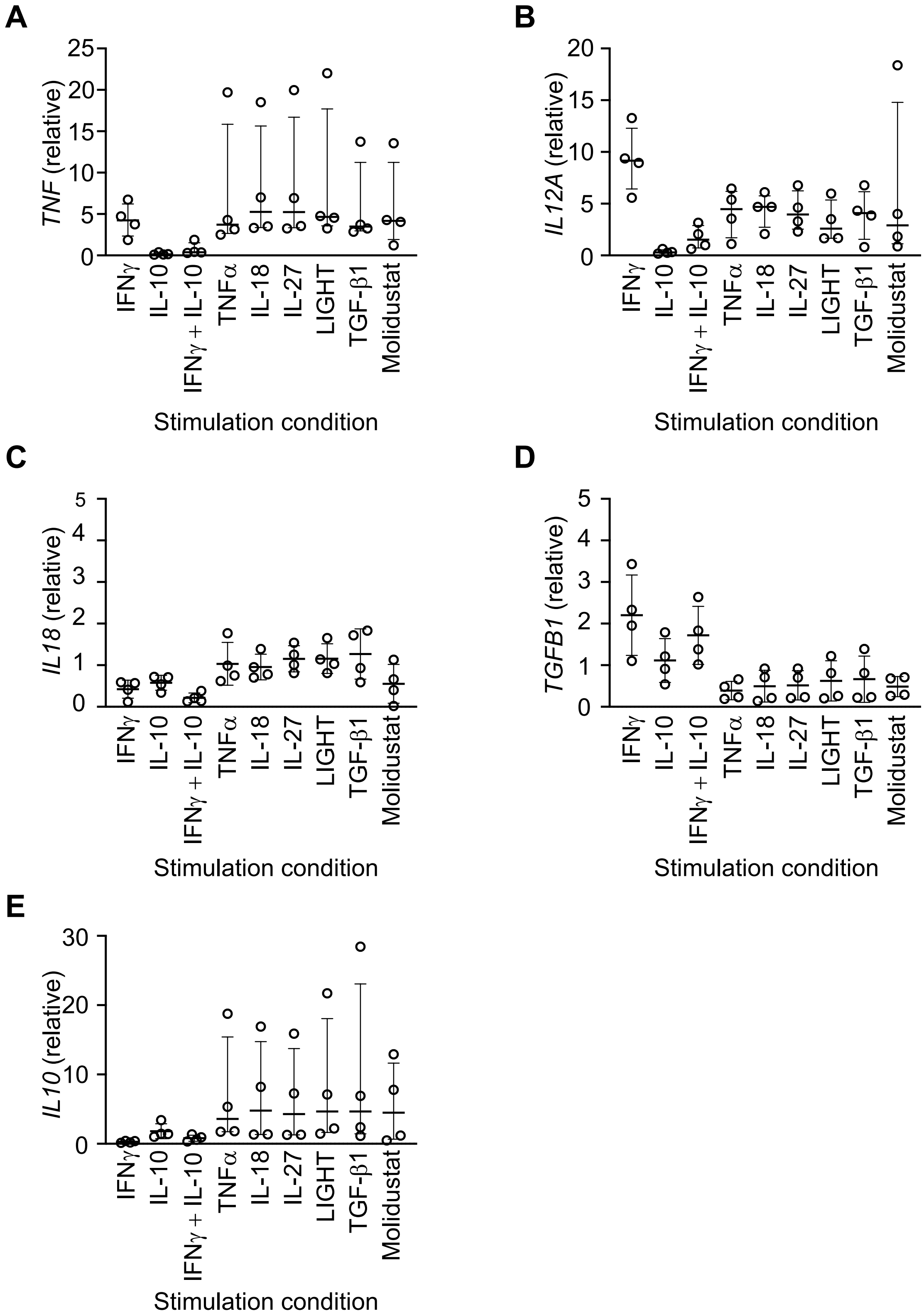


**Figure S6. IFNγ and IL-10 do not cooperate in stimulation of cytokine production by monocytes.** Peripheral blood monocytes from healthy subjects were treated ex vivo with the cytokines shown, and expression levels of cytokines were measured by qPCR. Gene expression levels were normalized to unstimulated condition for each individual.

Supplemental Table 1. Characteristics of participants in single-cell RNA-sequencing analysis

| Variables | Healthy subject (N = 3) | Favorable outcome (N = 4) | Unfavorable outcome (N = 7) |
|---|---|---|---|
| Age, Mean (SD), y | – | 65 ± 17 | 69 ± 7 |
| Male, N (%) | 1 (33) | 3 (75) | 2 (29) |
| Comorbidities, N (%) | | | |
| Diabetes | | 2 (50) | 2 (29) |
| Chronic immunosuppression | | 0 | 1 (14) |
| Cancer with chemotherapy | | 0 | 1 (14) |
| Hematologic malignancy | | 0 | 1 (14) |
| Current smoker | | 0 | 1 (14) |
| Arrest in public location, N (%) | | 3 (75) | 1 (14) |
| Witnessed arrest, N (%) | | 4 (100) | 4 (57) |
| Bystander performed CPR, N (%) | | 3 (75) | 4 (57) |
| Shockable rhythm, N (%) | | 2 (50) | 2 (29) |
| Time from collapse to ROSC, median (IQR), min | | 5 (2–7) | 10 (4–27) |
| Cardiac cause, N (%) | | 3 (75) | 5 (71) |
| TTM implementation, N (%) | | 2 (50) | 5 (71) |
| Pressor requirement, N (%) | | 2 (50) | 5 (71) |
| CPC at hospital discharge, N (%) | | | |
| 1 | | 4 (100) | |
| 2 | | 0 | |
| 3 | | | 2 (29) |
| 4 | | | 1 (14) |
| 5 | | | 4 (57) |

CPC, Cerebral Performance Category; CPR, cardiopulmonary resuscitation; ROSC, return of spontaneous circulation; TTM, target temperature management

Table S7. Characteristics of participants in flow cytometry analysis

| Variables | Healthy subject (N = 12) | Favorable outcome (N = 14) | Unfavorable outcome (N = 14) |
|---|---|---|---|
| Age, Mean (SD), y | – | 62 ± 16 | 68 ± 10 |
| Male, N (%) | 6 (50) | 10 (71) | 4 (29) |
| Arrest in public location, N (%) | | 13 (93) | 7 (50) |
| Witnessed arrest, N (%) | | 13 (93) | 9 (64) |
| Bystander performed CPR, N (%) | | 11 (79) | 8 (57) |
| Shockable rhythm, N (%) | | 9 (64) | 4 (29) |
| Time from collapse to ROSC, median (IQR), min | | 3 (2–10) | 15 (7–43) |
| Cardiac cause, N (%) | | 10 (71) | 6 (43) |
| CPC at hospital discharge, N (%) | | | |
| 1 | | 11 (79) | |
| 2 | | 3 (21) | |
| 3 | | | 4 (25) |
| 4 | | | 0 |
| 5 | | | 10 (71) |

CPC, Cerebral Performance Category; CPR, cardiopulmonary resuscitation; ROSC, return of spontaneous circulation

Table S8. Characteristics of participants in plasma cytokine/chemokine assay

| Variables | Healthy subject (N = 15) | Favorable outcome (N = 18) | Unfavorable outcome (N = 29) |
|---|---|---|---|
| Age, Mean (SD), y | 68 ± 12 | 59 ± 15 | 70 ± 13 |
| Male, N (%) | 8 (53) | 10 (56) | 13 (45) |
| Arrest in public location, N (%) | - | 15 (83) | 13 (45) |
| Witnessed arrest, N (%) | - | 15 (83) | 18 (62) |
| Bystander performed CPR, N (%) | - | 16 (89) | 18 (67) |
| Shockable rhythm, N (%) | - | 11 (65) | 11 (38) |
| Time from collapse to ROSC, median (IQR), min | - | 5.5 (2–13) | 20 (10–34) |
| CPC at hospital discharge, N (%) | - | | |
| 1 | | 16 (89) | - |
| 2 | | 2 (11) | - |
| 3 | | - | 3 (10) |
| 4 | | - | 0 |
| 5 | | - | 26 (90) |

CPC, Cerebral Performance Category; CPR, cardiopulmonary resuscitation; ROSC, return of spontaneous circulation
